# TLR7/8 stress response drives histiocytosis in SLC29A3 disorders

**DOI:** 10.1101/2022.10.27.513971

**Authors:** Takuma Shibata, Ryota Sato, Masato Taoka, Shin-Ichiroh Saitoh, Mayumi Komine, Kiyoshi Yamaguchi, Susumu Goyama, Yuji Motoi, Jiro Kitaura, Kumi Izawa, Yoshio Yamauchi, Yumiko Tsukamoto, Takeshi Ichinohe, Etsuko Fujita, Ryosuke Hiranuma, Ryutaro Fukui, Yoichi Furukawa, Toshio Kitamura, Toshiyuki Takai, Arinobu Tojo, Mamitaro Ohtsuki, Umeharu Ohto, Toshiyuki Shimizu, Manabu Ozawa, Nobuaki Yoshida, Toshiaki Isobe, Eicke Latz, Kojiro Mukai, Tomohiko Taguchi, Kensuke Miyake

**Affiliations:** Division of Innate Immunity, Department of Microbiology and Immunology, The Institute of Medical Science, The University of Tokyo, 4-6-1 Shirokanedai, Minato-ku, Tokyo 108-8639, Japan; Department of Chemistry, Graduate School of Science, Tokyo Metropolitan University, Minami-Osawa 1-1, Hachioji, Tokyo 192-0397, Japan; Department of Dermatology, Jichi Medical University, 3311-1 Yakushiji, Shimotsuke, Tochigi 329-0498, Japan; Division of Clinical Genome Research, The Institute of Medical Science, The University of Tokyo, Tokyo, Japan; Division of Molecular Oncology, Department of Computational Biology and Medical Sciences, Graduate School of Frontier Sciences, The University of Tokyo, Tokyo, 108-8639, Japan; Atopy Research Center, Juntendo University Graduate School of Medicine, Tokyo, 113-8421, Japan; Department of Mycobacteriology, Leprosy Research Center, National Institute of Infectious Diseases, Tokyo, 189-0002, Japan; Division of Viral Infection, Department of Infectious Disease Control, International Research Center for Infectious Diseases, The Institute of Medical Science, The University of Tokyo, Minato-ku, Tokyo 108-8639, Japan; Division of Cellular Therapy, The Institute of Medical Science, The University of Tokyo, Tokyo, 108-8639, Japan; Department of Experimental Immunology, Institute of Development, Aging and Cancer, Tohoku University, Sendai, Japan; Department of Hematology and Oncology, Research Hospital, The Institute of Medical Science, The University of Tokyo, Tokyo, Japan; Graduate School of Pharmaceutical Sciences, The University of Tokyo, 7-3-1 Hongo, Bunkyo-Ku, Tokyo 113-0033, Japan; Laboratory of Developmental Genetics, Center for Experimental Medicine and Systems Biology, The Institute of Medical Science, The University of Tokyo, 4-6-1 Shirokanedai, Minato-ku, Tokyo 108-8639, Japan; Institute of Innate Immunity, University Hospital Bonn, University of Bonn, Bonn, Germany; Laboratory of Organelle Pathophysiology, Department of Integrative Life Sciences, Graduate School of Life Sciences, Tohoku University, Sendai, Japan

## Abstract

SLC29A3, also known as ENT3, is a lysosomal transmembrane protein that transports nucleosides from the lysosomes to the cytoplasm^1^. Loss-of-function mutations in *SLC29A3* cause lysosomal nucleoside storage and histiocytosis: phagocyte accumulation in multiple organs^2,3^. However, little is known about the mechanism through which lysosomal nucleoside storage drives histiocytosis. Herein, histiocytosis in *Slc29a3*^−/−^ mice was demonstrated to depend on TLR7, which senses a combination of nucleosides and oligoribonucleotides^4,5^. TLR7 responded to lysosomal nucleoside storage and enhanced proliferation of Ly6C^hi^ CX3CR1^low^ immature monocytes and their maturation into Ly6C^low^ phagocytes in *Slc29a3*^−/−^ mice. Because accumulated nucleosides primarily originated from cell corpse phagocytosis, TLR7 in immature monocytes recognized nucleoside storage as lysosomal stress and increased phagocyte numbers. This non-inflammatory compensatory response is referred to as the TLR7 stress response where Syk, GSK3β, β-catenin, and mTORC1 serve as downstream signalling molecules. In SLC29A3 disorders, histiocytosis accompanies inflammation^6,7^. Nucleoside storage failed to induce pro-inflammatory cytokine production in *Slc29a3*^−/−^ mice, but enhanced ssRNA-dependent pro-inflammatory cytokine production in Ly6C^hi^ classical monocytes and peripheral macrophages, not proliferating immature monocytes. Patient-derived monocytes harbouring G208R *SLC29A3* mutation showed higher survival and proliferation in the presence of M-CSF and produced larger amounts of IL-6 upon ssRNA stimulation than did those derived from healthy subjects. A TLR8 antagonist inhibited the survival/proliferation of patient-derived macrophages. These results demonstrated that TLR7/8 responses to lysosomal nucleoside stress drive SLC29A3 disorders.

## Main

TLR7 and 8 are known as lysosomal ssRNA sensors that initiate innate immune responses during infections^8,9^. However, their structures indicate that they respond to RNA degradation products; TLR7 binds to guanosine (Guo) or deoxyguanosine (dGuo), as well as to uridine (Urd)-containing oligoribonucleotides (ORNs), whereas TLR8 interacts with Urd and purine-containing ORNs^4,5,10^. RNA degradation in endosomes/lysosomes proceeds to nucleosides, which are then transported to the cytoplasm for further degradation. SLC29A3, also known as ENT3, is a lysosomal nucleoside transporter abundantly expressed in macrophages^1^. Loss-of-function mutations in *SLC29A3* cause monogenic diseases, including H syndrome, Faisalabad histiocytosis, pigmented hypertrichosis with insulin-dependent diabetes mellitus syndrome, and familial Rosai-Dorfman disease^2,11,12^. These SLC29A3 disorders are characterised by histiocytosis: mononuclear phagocyte accumulation in multiple organs^2,11,12^. However, the mechanisms by which lysosomal nucleoside storage increases phagocyte numbers have not been elucidated. We hypothesised that the accumulated nucleosides act on TLR7 and TLR8 to drive histiocytosis in SLC29A3 disorders.

### TLR7-dependent histiocytosis in *Slc29a3*^−/−^ mice

*Slc29a3*^−/−^ mice were obtained (Extended Data Fig. 1a–1c), and various organs of these mice were examined by liquid chromatography-mass spectrometry (LC-MS) to evaluate nucleoside accumulation. We found significant increases in cytidine (Cyd), Guo, 2’-deoxycytidine (dCyd), dGuo, and thymidine (dThd) in the spleens of lysosomal nucleoside transporter-deficient *Slc29a3*^−/−^ mice (Fig. 1a, Extended Data Fig. 2). Significant accumulation of nucleosides was not observed in the other organs, such as the liver, kidney, heart, and lung.

**Fig. 1.**
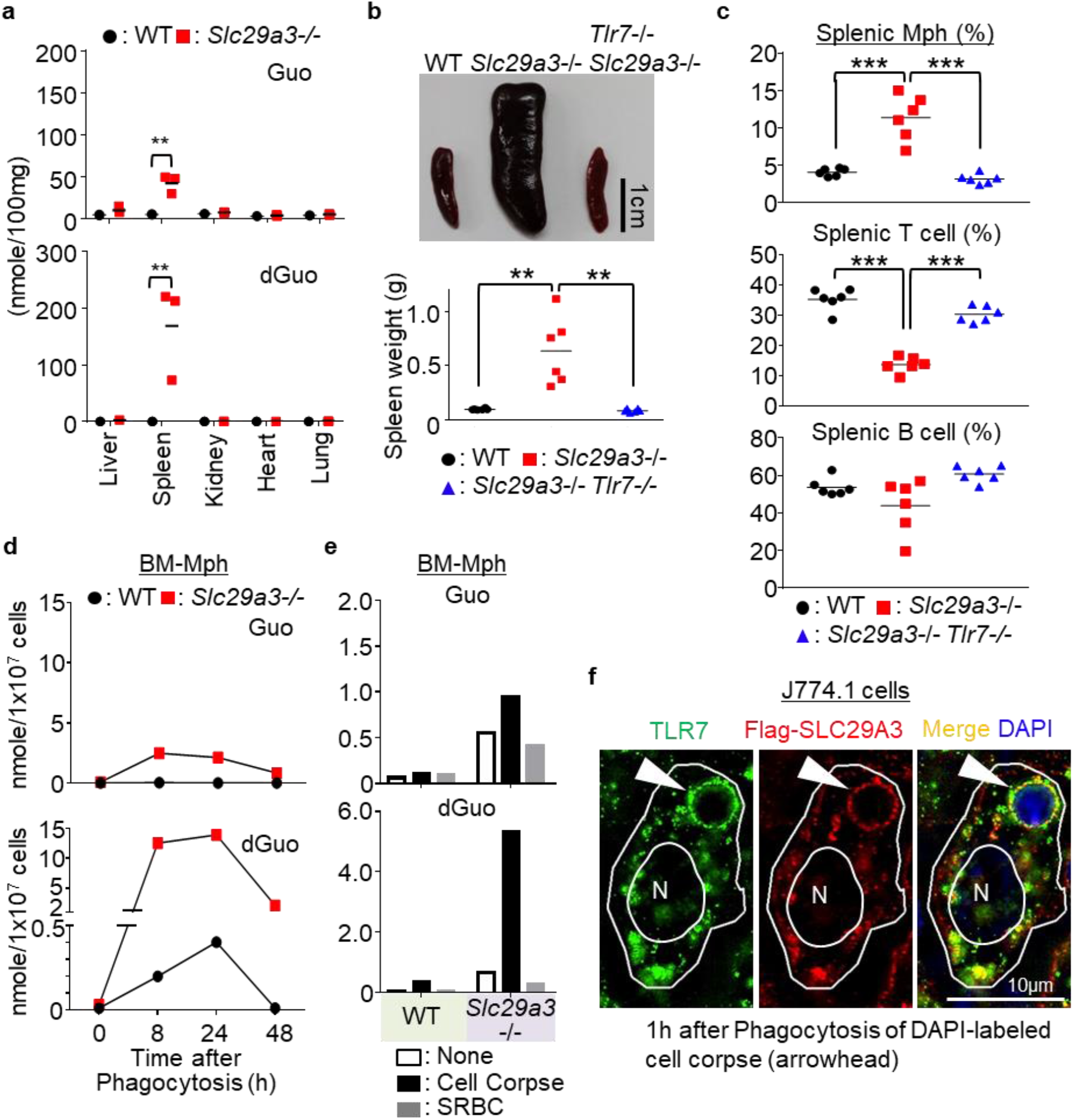
TLR7-dependent histiocytosis in *Slc29a3*^−/−^ mice. **a**, Amounts of guanosine (Guo) and 2’-deoxyguanosine (dGuo) in the indicated organs. Each dot represents a value (nanomole/100 mg tissue) from each mouse. **b**, Representative spleen images from 6-month-old mice (top). Scale bar, 1 cm. The bottom panel shows the spleen weight (n = 6). **c**, Percentages of NK1.1^-^ Ly6G^-^ CD11b^+^ monocytes, CD3ε^+^ T cells, and CD19^+^ B cells in the spleen from the indicated mice (n = 6). **d**, **e**, The amount of accumulated nucleosides (nanomoles) in 10^7^ cells of WT and *Slc29a3*^−/−^ BM-Mphs after treatment with 10^8^ dying thymocytes (cell corpse) or 10^9^ SRBCs for the indicated hours (**d**) or 24 h (**e**) were evaluated by LC-MS. **f**, Staining of TLR7 and Flag-SLC29A3 in the J774.1 macrophage cell line at 1 h after phagocytosis of the DAPI-labelled cell corpse. Arrowheads indicate a phagosome containing cell corpses. scale bar, 10 μm. \*\**p* < 0.01, \*\*\**p* < 0.001.

Because TLR7 responds to Guo and dGuo^4^, their accumulation in *Slc29a3*^−/−^ mice may activate TLR7. Therefore, we generated *Slc29a3*^−/−^*Tlr7*^−/−^ mice to evaluate the role of TLR7 in histiocytosis (Extended Data Fig. 1d–1f). Consistent with the previous report^3^, the spleens of *Slc29a3*^−/−^ mice were larger and heavier than those of wild-type (WT) mice due to increased cell number (Fig. 1b, Extended Data Fig. 3a). Concerning cell type-specific changes in the spleen and peripheral blood, increase in numbers was restricted to monocytes and macrophages, not T or B cells (Fig. 1c. Extended Data Fig. 3b-h). Peripheral blood platelet counts decreased, probably, due to premature clearance by accumulated macrophages (Extended Data Fig. 3i). All these changes were dependent on TLR7, as demonstrated in the *Slc29a3*^−/−^ *Tlr7*^−/−^ mice (Fig. 1b-c, Extended Data Fig. 3).

### Cell corpse phagocytosis increases nucleoside storage

Nucleoside storage was also observed in professional phagocytes, such as thioglycolate-elicited peritoneal macrophages (pMphs) and bone marrow-derived macrophages (BM-Mphs) (Extended Data Fig. 4a, 4b). In contrast, we observed much smaller nucleoside increases in splenic B cells and low nucleoside accumulation in BM-pDCs (Extended Data Fig. 4c, 4d). Given that B cells and BM-pDCs are less phagocytic than pMphs and BM-Mphs, phagocytosis might increase nucleoside storage. To address this possibility, we exposed dying thymocytes (cell corpse) to BM-Mphs (Fig. 1d, Extended Data Fig. 4e). We observed increases in nucleosides such as dGuo, dCyd, and dThd, which peaked at 8-24 h after cell corpse treatment. At 48 h, the levels of dGuo returned to normal in WT BM-Mphs but remained high in *Slc29a3*^−/−^ BM-Mphs. A lower, but appreciable, increase in the levels of Guo and Cyd was observed. In contrast to cell corpse phagocytosis, SRBC engulfment did not increase the amounts of nucleosides (Fig. 1e, Extended Data Fig. 4f). As SRBCs do not have nuclei, nuclear DNA and RNA from cell corpses are likely to be the major sources of nucleosides accumulated in BM-Mphs. Nucleoside storage in pMphs and BM-Mphs suggested their engulfment of cell corpses during elicitation by thioglycolate *in vivo* or by *in vitro* culture with M-CSF, respectively (Extended Data Fig. 4a, 4b).

We, next, studied the localisation of TLR7 and SLC29A3 in the mouse macrophage cell line J774.1, which engulfed the DAPI-labelled cell corpse. TLR7 and FLAG-tagged SLC29A3 were recruited to the cell corpse-containing phagosomes (Fig. 1f). These results suggest that SLC29A3 silences TLR7 by transporting phagosomal nucleosides to the cytoplasm in WT macrophages.

### TLR7 drives proliferation to increase phagocytes

Monocyte progenitors in the bone marrow give rise to Ly6C^hi^ monocytes/macrophages, which mature into Ly6C^low^ monocytes/macrophages^13^. In the spleen and peripheral blood, both Ly6C^hi^ and Ly6C^low^ monocytes increased in a TLR7-dependent manner in the *Slc29a3*^−/−^ mice (Fig. 2a, 2b, Extended Data Fig. 5a). Both Ly6C^hi^ and Ly6C^low^ splenic monocytes expressed TLR7 (Fig. 2c) and stored nucleosides, such as Guo and dGuo (Fig. 2d, Extended Data Fig. 5b), suggesting that TLR7 is cell-autonomously activated in these subsets. To characterise TLR7 responses in these monocyte subsets, we performed transcriptome analyses comparing *Slc29a3*^−/−^ and *Slc29a3*^−/−^ *Tlr7*^−/−^ monocytes with WT monocytes. An increase of 1.5-fold altered genes was observed in a TLR7-dependent manner in Ly6C^hi^ monocytes (Extended Data Fig. 5c). Gene set enrichment analyses (GSEAs) of more than 1.5-fold altered genes revealed that proliferation-related gene sets such as “E2F targets”, “G2M checkpoint”, and “mitotic spindle” were positively enriched in *Slc29a3*^−/−^ Ly6C^hi^ monocytes (Fig. 2e). These changes were dependent on TLR7 because such changes were not observed in *Slc29a3*^−/−^ *Tlr7*^−/−^ Ly6C^hi^ monocytes. To directly study the survival and proliferation of monocytes, splenic Ly6C^hi^ and Ly6C^low^ monocytes were sorted and cultured *in vitro* in the presence of M-CSF, which has been shown to promote histiocytosis in *Slc29a3*^−/−^ mice^3^. Ly6C^hi^, but not Ly6C^low^, monocytes from *Slc29a3*^−/−^ mice showed higher survival and proliferation in the presence of M-CSF at concentrations comparable to those *in vivo* (Fig. 2f). As the cell surface expression of the M-CSF receptor CD115 was not appreciably upregulated in *Slc29a3*^−/−^ splenic monocytes (Extended Data Fig. 5d), TLR7 did not increase cell surface CD115 to augment M-CSF response. The two signals via TLR7 or CD115 would synergistically drive the survival and proliferation of Ly6C^hi^ monocytes.

**Fig. 2.**
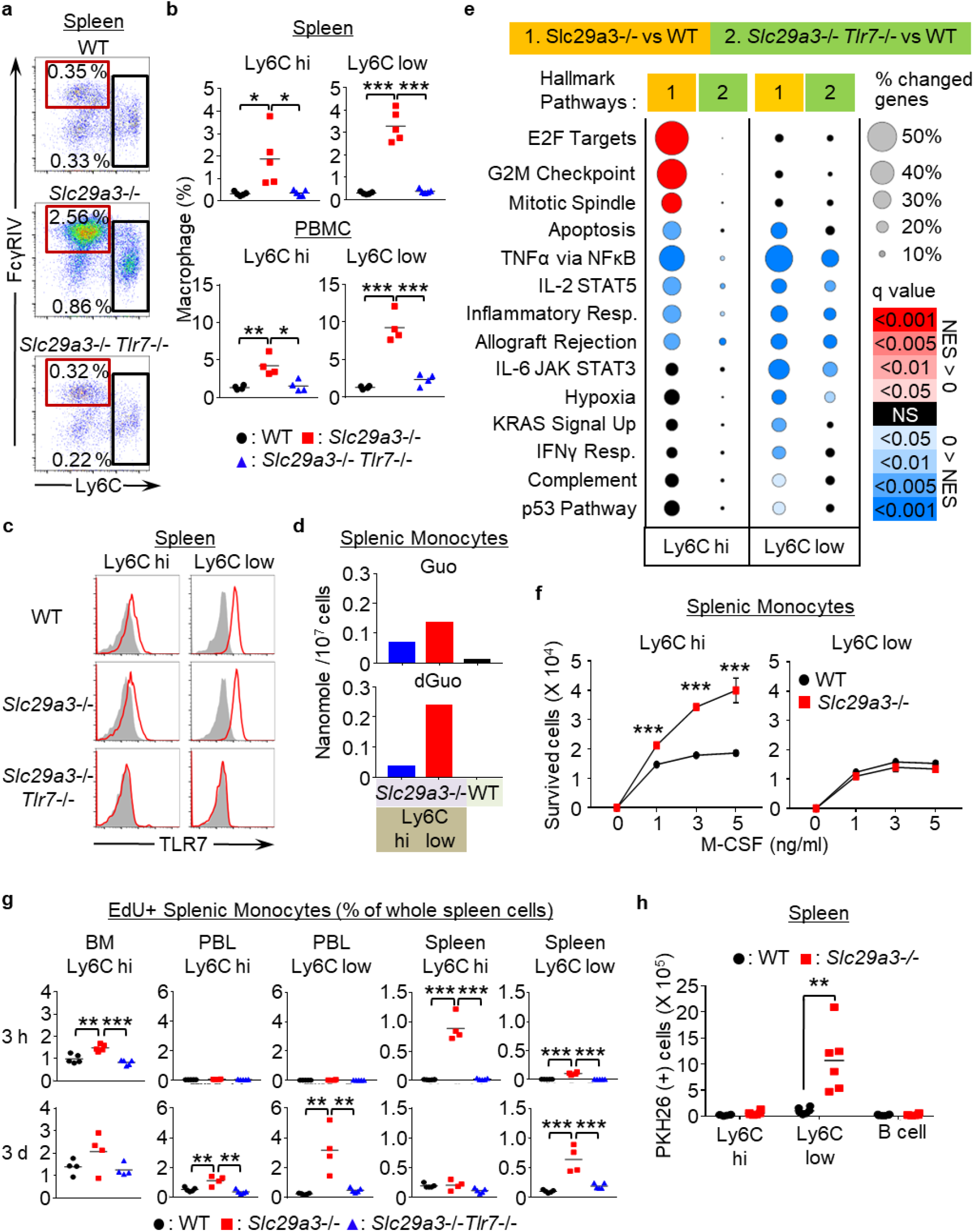
Ly6C^hi^ monocytes proliferate to increase Ly6C^low^ phagocytes. **a**, Representative FACS analyses of CD11b^+^ Ly6G^-^ NK1.1^-^ CD11C^low^ IA/IE^low^ splenic monocytes from WT, *Slc29a3*^−/−^, and *Slc29a3*^−/−^ *Tlr7*^−/−^ mice. The red and black squares show the gates of Ly6C^low^ and Ly6C^hi^ monocytes, respectively. **b**, Dot plots show the percentages of Ly6C^low^ and Ly6C^hi^ monocytes in the peripheral blood and spleen from the indicated mice (n = 5). **c,** Red histograms show intracellular TLR7 expression levels in Ly6C^hi^ and Ly6C^low^ monocytes from the indicated mice. Grey histograms show staining with the isotype control antibodies. **d**, Amounts (nmol/10^7^ cells) of Guo and dGuo in WT CD11b^+^ splenic monocytes or in Ly6C^hi^ and Ly6C^low^ splenic monocytes from *Slc29a3*^−/−^ mice. **e**, Gene set enrichment analysis (GSEA) of more than 1.5-fold changed genes in comparing Ly6C^hi^ and Ly6C^low^ splenic monocytes from *Slc29a3*^−/−^ and *Slc29a3*^−/−^*Tlr7*^−/−^ mice with those from WT mice. Red and blue circles indicate positive and negative normalised enrichment scores (NES), respectively. Their sizes indicate the percentage of genes with a >1.5-fold change in each gene set. The colour gradation indicates the *q*-value of positive/negative enrichment. **f**, Numbers of splenic Ly6C^hi^ and Ly6C^low^ monocytes from WT (black circle) and *Slc29a3*^−/−^ (red square) mice that survived for 4 days in *in vitro* culture with M-CSF at the indicated concentrations. Average values with s.d. from triplicate samples are shown. **g**, Uptake of the thymidine analogue EdU *in vivo* by monocytes from WT, *Slc29a3*^−/−^, and *Slc29a3*^−/−^*Tlr7*^−/−^mice at 3 h (upper) and 3 d (below) after intravenous EdU administration. Percentages of EdU^+^ cells in whole splenocytes are shown. n=5. \**p* < 0.05, \*\**p* < 0.01, \*\*\**p* < 0.001. **h**, Number of PKH26(+) cells in the indicated cell populations from the mice that had received PKH26-labelled dying thymocytes 1 h before analyses. Each dot represents the value for each mouse (n = 6).

To study the *in vivo* proliferation of monocytes, mice were intravenously administered with the thymidine analogue–EdU, and the percentages of EdU^+^ monocytes in bone marrow, peripheral blood, and spleen were analysed 3-h post EdU administration. In WT mice, Edu^+^ proliferating monocytes were found only in bone marrow Ly6C^hi^ monocytes (Fig. 2g). The percentage of Edu^+^ monocytes in the bone marrow TLR7-dependently increased in *Slc29a3*^−/−^mice. Even more drastic changes were observed in the spleen, where Edu^+^ Ly6C^hi^ monocytes were found only in *Slc29a3*^−/−^ mice. We analysed EdU^+^ monocytes 3 days after EdU administration and observed that the majority of Edu^+^ monocytes in the circulation and spleen turned Ly6C^low^ (Fig. 2g), suggesting that proliferating Ly6C^hi^ monocytes mature into Ly6C^low^ monocytes within 3 days in the *Slc29a3*^−/−^ mice. Ly6C^low^ monocytes in the *Slc29a3^−/−^* mice stored deoxyribonucleosides more than Ly6C^hi^ monocytes (Fig. 2d, Extended Data Fig. 5b). Because lysosomal deoxyribonucleosides were derived from cell corpses (Fig. 1d, 1e), Ly6C^low^ monocytes are likely to have engulfed cell corpses during or after maturation from Ly6C^hi^ monocytes. Consistent with this, splenic Ly6C^low^ monocytes in the *Slc29a3*^−/−^ mice engulfed intravenously administered dying thymocytes (Fig. 2h). TLR7, therefore, increased the number of phagocytes in the *Slc29a3*^−/−^ mice. TLR7 is likely to recognise lysosomal nucleoside storage as lysosomal stress and increases phagocyte number as a compensatory mechanism.

### Mutually exclusive induction of proliferation and inflammation by TLR7

To further narrow down the proliferating population of Ly6C^hi^ monocytes, we examined a marker specifically expressed or not expressed in proliferating Ly6C^hi^ monocytes. The cell surface expression of CX3CR1, a chemokine receptor for the membrane-tethered chemokine CX3CL1^14^, increases with maturation from Ly6C^hi^ to Ly6C^low^ monocytes^15^. We found that the CX3CR1^low^ population in Ly6C^hi^ monocytes increased in a TLR7-dependent manner in the *Slc29a3*^−/−^ mice (Fig. 3a). When splenic monocytes were cultured *in vitro* for 1 h with EdU, its uptake was detected in this immature monocyte subset but not in more mature Ly6C^hi^ CX3CR1^hi^ classical monocytes (Fig. 3b). These results demonstrate that Ly6C^hi^ CX3CR1^low^ immature monocytes proliferate in response to lysosomal nucleoside storage in the *Slc29a3*^−/−^ mice.

**Fig. 3.**
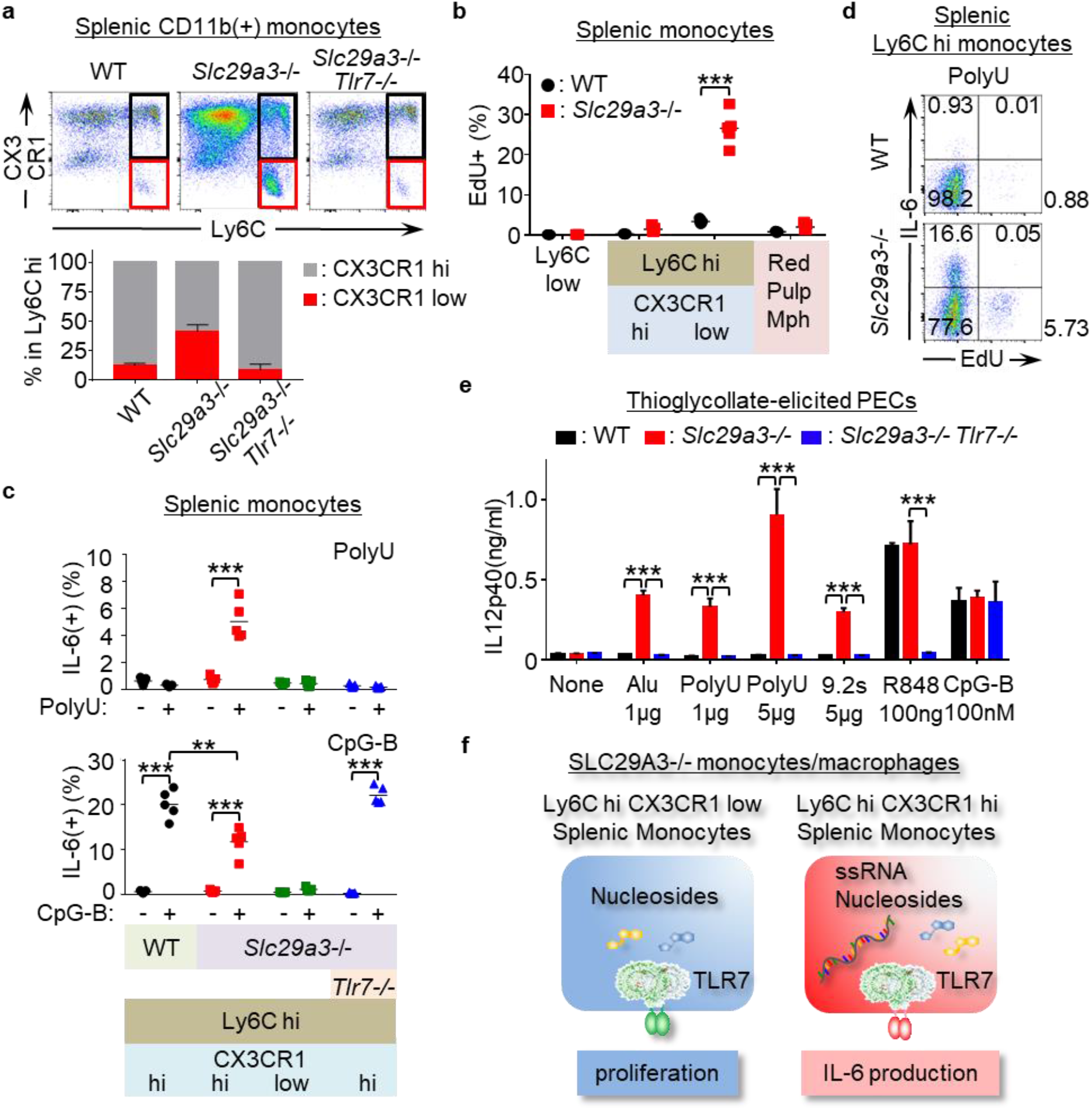
TLR7 differentially induces proliferation and inflammation. **a**, Expression of CX3CR1 and Ly6C in splenic CD11b^+^ Ly6G^-^ NK1.1^-^ CD11C^low^ IA/IE^low^ monocytes from WT, *Slc29a3*^−/−^, and *Slc29a3*^−/−^ *Tlr7*^−/−^ mice (top). Black and red gates show Ly6C^hi^ CX3CR1^hi^ and Ly6C^hi^ CX3CR1^low^ monocytes, respectively. The bottom panel shows the percentages of the two Ly6C^hi^ monocyte subsets in the indicated mice (n = 4). **b**, EdU uptake by each splenic monocyte subset during 1 h culture with 10 μM EdU. Each dot represents a value from a single mouse (n = 4). **c**, Percentage of IL-6^+^ cells in each monocyte subset after *in vitro* stimulation with poly U (10 μg/mL) or CpG-B (1 μM) for 4 h. Brefeldin A (10 μg/mL) was added during cell stimulation. Each dot shows the values for each mouse from the indicated mice (n = 5). **d**, Representative dot plot of EdU^+^ and IL-6^+^ cells in Ly6C^hi^ splenic monocytes treated with poly U + brefeldin A for 4 h. EdU was added during the last hour of stimulation. **e**, IL-12 p40 production by thioglycolate-elicited pMphs after stimulation with TLR7 and TLR9 ligands for 18 h. Alu: Alu retroelements. The results are represented as the average with s.d. from triplicate samples. **f**, Schematics showing the induction of two distinct TLR7 responses, proliferation and cytokine production, in *Slc29a3*^−/−^ mice. \*\**p* < 0.01, \*\*\**p* < 0.001.

Because SLC29A3 disorders are considered to be inflammatory diseases in which IL-6 has pathogenic roles^6^, we examined TLR7-dependent inflammatory responses in the *Slc29a3*^−/−^ mice. Unexpectedly, inflammation-associated gene sets such as “TNFα via NF-κB” and “inflammatory response” were negatively enriched in splenic Ly6C^hi^ monocytes (Fig. 2e). Consistent with this, SLC29A3-deficiency in Ly6C^hi^ monocytes did not increase the expression levels of mRNAs encoding proinflammatory cytokines, such as IFN-α, IFN-β, IFN-γ, IL-6, IL-17A, IL-23, and TNF-α (Extended Data Fig. 6a). Furthermore, proinflammatory cytokines, such as TNF-α, IFN-β, IL-1β, IFN-α, and IL-6, were not identifiable in the serum using ELISA (Extended Data Fig. 6b). These results suggest that nucleoside accumulation is not sufficient to induce TLR7-dependent inflammation. We previously reported that TLR7 response to ssRNA is enhanced by Guo/dGuo^4^, which accumulated in *Slc29a3*^−/−^ monocytes (Fig. 2d). Therefore, TLR7 response to ssRNA may be enhanced in *Slc29a3*^−/−^ monocytes. As predicted, *Slc29a3*^−/−^ Ly6C^hi^ CX3CR1^hi^ classical monocytes produced IL-6 upon poly U stimulation, whereas WT Ly6C^hi^ macrophages did not respond to poly U (Fig. 3c). In contrast, *Slc29a3*^−/−^ Ly6C^hi^ CX3CR1^low^ proliferating immature monocytes did not respond to poly U treatment (Fig. 3c). Consistent with this finding, when splenic monocytes were treated with EdU and poly U, we only detected EdU^+^ or IL-6^+^Ly6C^hi^ monocytes but not EdU^+^ IL-6^+^ double-positive monocytes (Fig. 3d). TLR7 is likely to induce proliferation and inflammation in a mutually exclusive manner in immature and classical monocytes, respectively.

*Slc29a3*^−/−^ Ly6C^hi^ classical monocytes responded to the TLR9 ligand CpG-B but produced smaller amounts of IL-6 than WT Ly6C^hi^ classical monocytes (Fig. 3c). In addition, *Slc29a3*^−/−^ Ly6C^low^ monocytes did not produce IL-6 in response to CpG-B, whereas WT Ly6C^low^ monocytes did (Extended Data Fig. 7a). As TLR7-deficiency rescued impaired TLR9 response (Fig. 3c, Extended Data Fig. 7a), nucleoside-activated TLR7 is likely to inhibit TLR9 responses.

An antibody array for cytokines showed that poly U-stimulated splenic monocytes produced chemokines and cytokines, including CCL2, CCL3, CCL12, CXCL2, CXCL9, IL-6, TNF-α, IL-12p40, and IL-10, in *Slc29a3*^−/−^ mice (Extended Data Fig. 7b, 7c). Furthermore, *Slc29a3*^−/−^ professional macrophages, such as BM-Mphs and pMphs, exhibited stronger IL-12p40 production in response to poly U than did WT macrophages, and their responses to R848 and CpG-B were not impaired (Fig. 3e, Extended Data Fig. 7d). Inflammation in SLC29A3 disorders might be driven by an enhanced TLR7 response to ssRNA in Ly6C^hi^ splenic monocytes and peripheral macrophages. These results suggest that the TLR7 response to nucleoside storage varies with monocyte maturation from proliferation to an excessive inflammatory response to ssRNA (Fig. 3f).

### Signals involved in TLR7-dependent monocyte proliferation

Next, we focused on growth-promoting TLR7 signalling. We cultured whole splenocytes *in vitro* with 3 ng/ml M-CSF, where larger numbers of *Slc29a3*^−/−^ splenic monocytes survived than WT monocytes (Fig. 2f). Inhibitors of MEK (PD0325901), Syk (PRT062607, R788), β-catenin (PKF118-120), PI3K (wortmannin), AKT (Afuresertib), mTORC1 (rapamycin, Torin1), and MyD88 (ST2825), but not JNK (JNK-IN-8), reduced the number of surviving monocytes (Fig. 4a). We examined the *in vivo* TLR7-dependent activation of these signalling molecules. Flow cytometry analyses of Ly6C^hi^ monocytes revealed TLR7-dependent increases in the activated form of β-catenin and the phosphorylated forms of signalling molecules including Syk, GSK3β, and the ribosomal protein S6 (the mTORC1 downstream effector) (Fig. 4b). In contrast, TLR7-dependent ERK phosphorylation was not significantly observed. These results suggest that TLR7 activates signalling molecules such as Syk, GSK3β, β-catenin, and mTORC1 to promote proliferation/survival in *Slc29a3*^−/−^ Ly6C^hi^ monocytes.

**Fig. 4.**
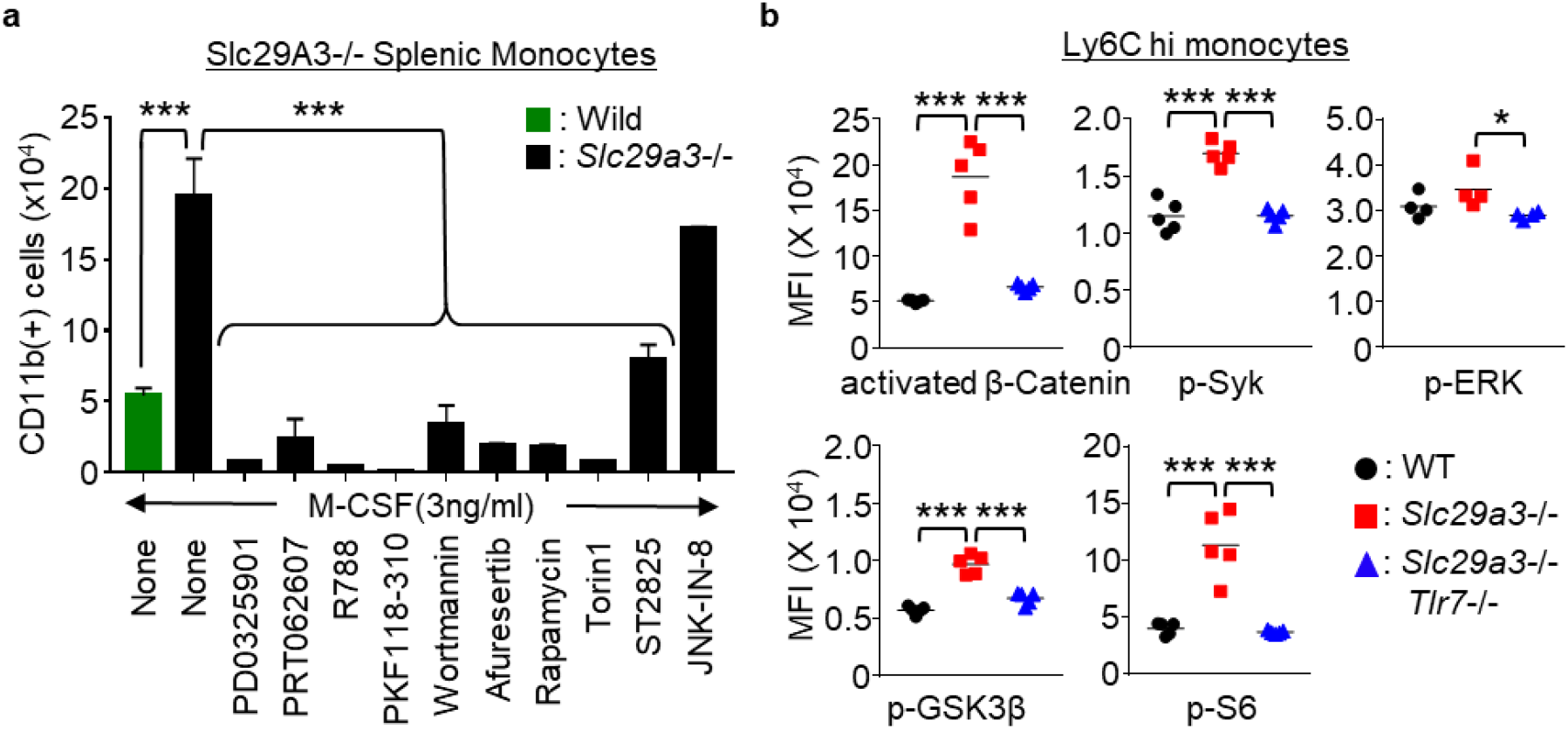
FcRγ and DAP10 mediate TLR7 responses in *Slc29a3*^−/−^ mice. **a**, Number of splenic CD11b^+^ macrophages from WT (green) or *Slc29a3*^−/−^(black) mice that survived *in vitro* 4 day culture in the presence of 3 ng/mL M-CSF under serum-free conditions. Cells were treated with inhibitors of MEK (PD0325901, 1 μM), Syk (PRT062007, 1 μM; R788, 0.5 μM), β-catenin (PKF118-120, 5 μM), PI3K (wortmannin, 10 μM), AKT (Afuresertib, 5 μM), mTORC1 (rapamycin, 0.5 μM), mTORC1 and 2 (Torin1, 250 nM), MyD88 (ST2825, 10 μM), and JNK (JNK-IN8, 1 μM). Bar graphs represent mean values ±s.d. of triplicate samples. **b**, Mean fluorescence intensity (MFI) of staining with activation- and phospho-specific Abs to signalling molecules in Ly6C^hi^ monocytes from the indicated mice (n = 4–5). \**p* < 0.05, \*\*\**p* < 0.001.

### TLR8-dependent histiocytosis in humans

Finally, we investigated whether human monocytes from SLC29A3 disorders show enhanced TLR7/8 responses. In human peripheral blood mononuclear cells (PBMCs) from a patient harbouring the *SLC29A3* p.Gly208Arg (G208R) mutation^20^, CD14^low^ CD16^hi^ monocytes, which are equivalent to the mouse Ly6C^low^ monocytes^13,21^, were increased by approximately 3-folds (Fig. 5a). The expression levels of TLR7 and TLR8 in both CD14^hi^ CD16^low^ and CD14^low^ CD16^hi^ monocytes were not altered in the patient (Fig. 5b). To study monocyte proliferation and survival, PBMCs were cultured *in vitro* with human M-CSF. A larger number of HLA-DR^+^ CD11b^+^ monocytes from the patient survived *in vitro* culture than those from three healthy subjects (Fig. 5c). PBMCs were allowed to differentiate into macrophages by human M-CSF and IL-4 and were stimulated with the TLR7/8 ligands poly U and RNA9.2S or TLR8 ligand ssRNA40^4^. Macrophages harbouring the G208R *SLC29A3* mutation produced larger amounts of IL-6 than the control macrophages in response to TLR7/8 and TLR8 ssRNA ligands, strongly suggesting that the *SLC29A3* mutation enhanced TLR8 responses in macrophages (Fig. 5d). We also examined PBMCs from another patient harbouring the *SLC29A3* p.Ser184Arg (S184R) mutation^22^. The percentage of monocytes did not increase (Extended Data Fig. 8a); however, proliferation and survival *in vitro* in the presence of M-CSF were significantly enhanced (Extended Data Fig. 8b). These results demonstrate that the phenotypes in the *Slc29a3*^−/−^ mice were consistent with those of the patients with SLC29A3 mutation.

**Fig. 5.**
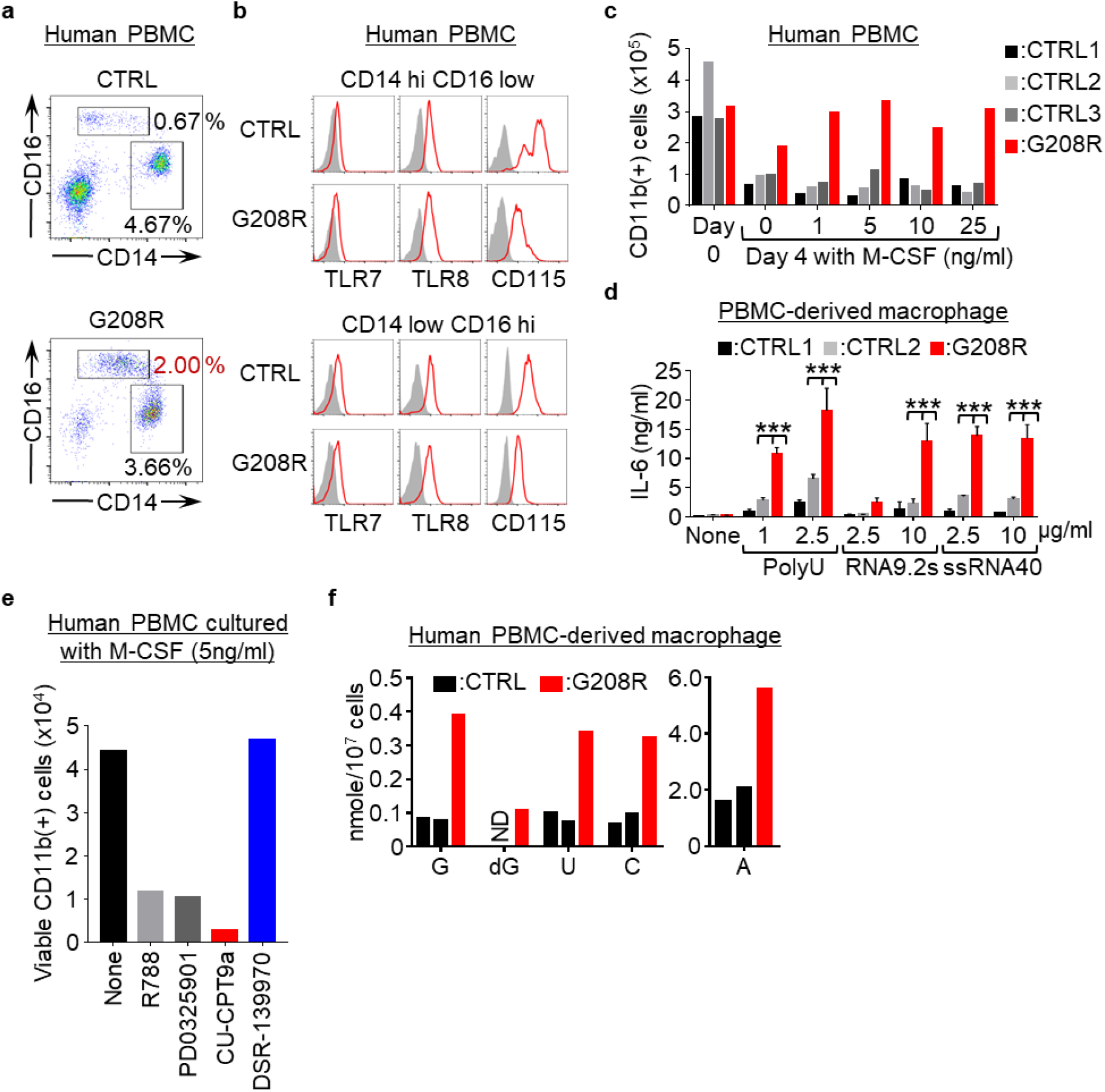
TLR8 drives histiocytosis in SLC29A3 disorders. **a**, **b**, Expression of surface CD16/CD14/CD115 (**a**, **b**) and intracellular TLR7/TLR8 (**b**) in HLA-DR^+^ CD15^-^ CD56^-^ PBMCs from a patient with the G208R *SLC29A3* mutation and a healthy subject. Red and grey histograms show staining with the indicated and isotype-matched control antibodies, respectively. **c**, Number of surviving CD11b^+^ CD15^-^ CD56^-^ monocytes in PBMCs from patients (red) or healthy subjects (black and grey) after 4 days of culture with M-CSF at the indicated concentrations. **d**, PBMC-derived macrophages from the patient (red) or two control subjects (black and grey) were stimulated with the indicated TLR7 or TLR8 ssRNA ligands at the indicated concentrations. IL-6 production was evaluated using ELISA. The bars represent the average values with s.d. from triplicate samples. **e**, CD11b^+^ cells that survived *in vitro* PBMC culture with 5 ng/mL M-CSF in the presence of the indicated inhibitors of Syk (1 μM R788), MEK1/2 (1 μM PD0325901), TLR8 (10 μM CPT9a), and TLR7 (10 μM DSR-139970). **f**, The amounts (nmol/10^7^ cells) of nucleosides accumulated in PBMC-derived macrophages. ND, not detectable. \*\**p* < 0.01, \*\*\**p* < 0.001.

To determine whether TLR7 or TLR8 drives histiocytosis in humans, PBMCs from a patient harbouring the G208R or the S184R SLC29A3 mutations were cultured with human M-CSF in the presence of TLR7 or TLR8 specific inhibitors. The TLR8 antagonist CU-CPT9a^23^, but not the TLR7 antagonist DSR-139970 (Cpd-7)^24^, inhibited the *in vitro* survival of human macrophages with SLC29A3 mutation as much as the Syk inhibitors R788 and PD0325901 (Fig. 5e, Extended Data Fig.8b). Additionally, we detected approximately 3-fold higher amounts of Urd, the TLR8 ligand^4,8,10^, in macrophages harbouring the G208R *SLC29A3* mutation than in those from healthy subjects (Fig. 5f). These results suggest that SLC29A3 negatively regulates TLR8 by transporting Urd from the lysosomes to the cytoplasm, and that TLR8 drives histiocytosis in SLC29A3 disorders. The role of huTLR7 in monocyte proliferation in SLC29A3 disorders remains unclear.

Here, we identified a previously unknown histiocytosis-driving program activated by germline loss-of-function mutations in *SLC29A3*. This program recognises lysosomal nucleoside storage as lysosomal stress and increases phagocytes by driving monocyte proliferation and maturation. This program is reminiscent of another response activated by haem following erythrocyte clearance. Although a haem sensor has not been identified, haem stress in macrophages induces their differentiation into red pulp macrophages, which are phagocytes specialised for red blood cell clearance^25^. We refer to metabolite-dependent macrophage proliferation and maturation as the lysosomal stress response. TLR7 and TLR8 serve as nucleoside sensors to activate lysosomal stress responses, which drive histiocytosis, unless the stress is relieved. These results demonstrate that SLC29A3 disorders are lysosomal stress diseases.

## Methods

### Generation of *Slc29a3*^−/−^ and *Tlr7*^−/−^ Mice

C57BL/6 *Slc29a3*^−/−^ and *Tlr7*^−/−^ mice were generated using the CRISPR/CAS9 system. gRNA target sites on *Slc29a3* and *Tlr7* were determined using CRISPRdirect software (http://crispr.dbcls.jp/), and 5’-agcttcttgatggttactcg-3’ and 5’-gaacagttggccaatctctc-3’ sequences were chosen for the construction of *Slc29a3*^−/−^ and *Tlr7*^−/−^ mice, respectively. These gRNA target sequences were cloned into the BbsI site of the pKLV-U6 gRNA (BbsI)-PGKpuro2ABFP vector (Addgene plasmid 50946). Using the constructed vectors as templates, gRNAs were synthesised by *in vitro* transcription using the MEGA shortscript T7 Transcription Kit (Thermo Fisher Scientific). Additionally, hCAS9 mRNA was synthesised from the hCAS9 sequence in pX458 (Addgene plasmid 48138) *in vitro* using the mMESSAGE mMACHIN T7 ULTRA Transcription Kit (Thermo Fisher Scientific, USA). To generate Slc29a3^−/−^ Tlr7^−/−^ mice, synthesized gRNAs (50 ng/μL) targeting the *Slc29a3* and *Tlr7* genes, hCAS9 mRNA (100 ng/μL), and the repair donor DNA (100 ng/μL) to introduce a stop codon into the Slc29a3 *locus* (5’-tgcctctgaggacaatgtataccacagctccaatgctgtctacagagcccTGATAGCGTAAAGCACTGAGGAAGcgagtaaccatcaagaagctgaccaggaagccctgctggggaaactacta-3’; Lower-case letters and capital letters represent homology arm and insertion sequence, respectively) were injected into zygotes from C57BL/6 mice at the pronuclei stage. The injected zygotes were then transferred to the oviducts of pseudopregnant female C57BL/6 mice. Candidates for *Slc29a3*^−/−^ *Tlr7*^−/−^ mice were typed by PCR using primer pairs for *Slc29a3* (Fw:5’-CCAGCATGGACGAGAGATGTCTTC-3’, Rv:5’-GCACCATTGAAGCGATCCTCTGG-3’) and *Tlr7* (Fw:5’-GAGGGTATGCCGCCAAATCTAAAGAATC-3’, Rv:5’-CTGATGTCTAGATAGCGCAATTGC-3’). The PCR product was analysed using the MCE-202 MultiNA Microchip Electrophoresis System for DNA/RNA Analysis (SHIMAZU, Japan), and *Slc29a3*^−/−^ *Tlr7*^−/−^ male mice were chosen for mating with WT C57BL/6 female mice. *Slc29a3*^+/−^ *Tlr7*^+/−^ mice were mated to establish *Slc29a3*^−/−^ and *Slc29a3*^−/−^ *Tlr7*^−/−^ mice. The sequences of the mutated allele in *Slc29a3*^−/−^ and *Tlr7*^−/−^ mice were confirmed by direct sequencing performed by FASMAC (Japan).

### Mice

WT C57BL/6 mice were purchased from Japan SLC, Inc. (Shizuoka, Japan). All animals were housed in SPF facilities at the Institute of Medical Science, University of Tokyo (IMSUT). All animal experiments were approved by the Institutional Animal Care and Use Committee of the IMSUT.

### Reagents

PD0325901 (mirdametinib), PRT062607 (P505-15) HCl, R788 (fostamatinib), wortmannin (KY 12420), afuresertib (GSK2110183), AY-22989 (Rapamycin), and JNK-IN-8 were purchased from Selleck Chemical (USA). PKF118-310 was purchased from Sigma (USA). Torin 1 and ST2825 were purchased from Calbiochem/Merck Millipore (USA) and ChemScence (USA), respectively.

Stable isotope-labelled nucleosides, G (13C10, 98%;15N5, 96-98%), U (13C9, 98%;15N2, 96-98%), A (13C10, 98%;15N5, 96-98%), C (13C9, 98%;15N3, 96-98%), and dG (13C10, 98%;15N5, 96-98%) were purchased from Cambridge Isotope Laboratories (Massachusetts, USA) for the quantification of nucleosides by LC-MS/MS. The EdU used in the *in vivo* proliferation assay was purchased from Tokyo Chemical Industry Co. (Tokyo, Japan).

DSR-139970 (Cpd7), a TLR7 inhibitor, was kindly provided by Sumitomo Pharma Co., Ltd. (Japan). CU-CPT9a, a specific TLR8 inhibitor, and R848 were purchased from InvivoGen (Hong Kong, China). Sheep red blood cells (SRBC) used in the phagocytic assay were obtained from Cosmo Bio Co. (Japan).

RNA9.2s (20mer, UsGsUsCsCsUsUsCsAsAsUsGsUsCsCsUsUsCsAsA), ssRNA40 (GsCsCsCsGsUsCsUsGsUsUsGsUsGsUsGsAsCsUsC), PolyU (19mer, UsUsUsUsUsUsUsUsUsUsUsUsUsUsUsUsUsUsU), and ODN1668 (20mer, dTsdCsdCsdAsdTsdGsdAsdCsdGsdTsdTsdCsdCsTdsdGsdAsdTsdGsdCsdT), in which ‘s’ depicts a phosphothioate linkage, were synthesized by FASMAC (Japan).

### Establishment of anti-human TLR7/8 monoclonal antibodies

To establish an anti-human TLR7/8 monoclonal antibody (mAb), WT Wistar rats and BALB/c mice were immunised several times with purified huTLR7/8 ectodomain and Ba/F3 cells expressing huTLR7/8 mixed with TiterMax Gold. Four days after final immunisation, splenocytes and SP2/O myeloma cells were fused with polyethylene glycol. After selection by hypoxanthine/aminopterin/thymidine (HAT), antibodies to TLR7 and TLR8were selected by flow cytometry analyses using Ba/F3 cells expressing huTLR7/8. Anti-huTLR7 and huTLR8 mAbs were designated rE3 (rat IgG2a/κ) and M7B (mouse IgG1/κ), respectively. The purity of the mAbs was checked by SDS-PAGE and Coomassie brilliant blue staining, and biotinylated mAbs were used for subsequent experiments.

### Flow cytometry

For the preparation of samples for flow cytometry analysis, the spleens were minced using glass slides, and the bone marrow cells were pipetted several times to disperse the cells in RPMI1640 culture medium. The suspended samples were teased using nylon mesh to remove tissue debris. All samples were treated with BD Pharm lysing buffer (BD Biosciences) to remove red blood cells before being subjected to cell staining. Cell surface staining for flow cytometry analysis was performed using fluorescence-activated cell sorting (FACS) staining buffer (1×PBS with 2.5% FBS and 0.1% NaN3). The prepared cell samples were incubated for 10 min with an unconjugated anti-mouse CD16/32 blocking mAb (clone 95) to prevent nonspecific staining in the staining buffer. The cell samples were then stained with fluorescein-conjugated mAbs for 15 min on ice.

Peripheral blood mononuclear cells (PBMCs), bone marrow cells (BMs), and splenocytes from mice were stained with fluorescent dye-conjugated monoclonal antibodies specific for the following markers after blocking antibody treatment: CD11b (clone M1/70), FcγRIV (clone 9E9), CD3ε (clone 145-2c11), CD19 (clone 6D5), CD11c (clone N418), CD71 (clone R17217), Ter119/Erythroid Cells (clone Ter119), Ly6C (clone HK1.4), and Ly6G (clone 1A8).

Human PBMCs were assessed using fluorescent dye-conjugated antibodies specific for the following markers: HLA-DR (clone G46-6), CD14 (clone M5E2), CD16 (clone 3G8), CD15 (clone HI98), CD56 (clone 5.1H11), CD3ε (clone SK7), and CD19 (clone HIB19). These antibodies were purchased from eBioscience (USA), BioLegend (USA), BD Biosciences (USA), and TONBO Biosciences (USA). Biotinylated anti-mouse TLR7 (A94B10) mAb was previously established in our laboratory^29^.

To detect endolysosomal TLR7 and TLR8, cells after cell surface staining were fixed and permeabilised using a Fixation/Permeabilization Solution Kit (BD Biosciences, USA) and stained again with biotinylated anti-mouse/human TLR7/8 mAb and PE streptavidin (Biolegend, USA). To detect intracellular mouse IL-6 in splenocytes stimulated with various ligands in the presence of Brefeldin A (10μg/ml), cells after cell surface staining were fixed with Fixation Buffer (Biolegend, USA), permeabilised using 1X Click-iT saponin-based permeabilisation and wash reagent (Invitrogen™ ThermoFisher Scientific, USA), and then stained with PE-conjugated anti-mouse IL-6 mAb (clone MP5-20F3, Biolegend). Stained cells were analysed using a BD LSR Fortessa cell analyser (BD Biosciences, USA) or an ID7000 Spectral Cell Analyser (Sony Biotechnology, Japan). All flow data were analysed using FlowJo software v10.7 (BD Biosciences, USA).

### Intracellular phospho-flow cytometry

Phosphorylation of signalling molecules was detected by flow cytometry. First, 3-4 × 10^6^ splenocytes after cell surface staining were fixed with Cyto-Fast Fix/Perm buffer (BioLegend, USA) for 20 min at room temperature and washed twice with FACS staining buffer. Fixed cells were further permeabilised by adding pre-chilled True-Phos Perm Buffer (BioLegend, USA) and incubated at −20 °C for 2–3 h. After washing twice with FACS staining buffer, permeabilised cells were stained with PE-conjugated anti-p-Syk (clone C87C1; 1:50 dilution; Cell Signaling Technology, Danvers, MA, USA), anti-p-S6 (clone D57.2.2E; 1:100 dilution; Cell Signalling Technology), anti-p-GSK3β (clone D85E12; 1:50 dilution; Cell Signalling Technology), anti-p44/42 MAPK (Erk1/2) (clone 137F5; 1:50 dilution; Cell Signalling Technology), or Fluor647-conjugated anti-active β-catenin (clone 8E7; 1:400 dilution; Merck Millipore) and subjected to flow cytometry analyses.

### Cell sorting of pDCs and splenic monocytes

Cell staining for sorting was performed in a sorting buffer (1×PBS with 10% FBS, 10 mM HEPES and 1 mM Sodium pyruvate). Flt3L induced bone marrow cells were stained with anti-CD11c/B220 mAbs and CD11c+B220+ cells were sorted as bone marrow-derived pDCs. For purification of Ly6Clow and Ly6Chigh splenic monocytes, whole splenocytes from wild-type and *Slc29a3*^−/−^ mice were sequentially incubated with biotinylated anti-mouse CD3 (clone 145-2C11)/CD19 (clone 6D5)/NK1.1 (clone PK136)/Ly6G (clone aA8)/TER-119/erythroid cells (clone Ter-119) and Streptavidin MicroBeads (Miltenyi Biotec, Germany). The magnetically labelled cells were removed using autoMACS (Miltenyi Biotec, Germany), and the enriched cells were stained with anti-mouse CD11b/Ly6C/Fcgr4/NK1.1/Ly6G/Siglec-F mAb. Ly6C^low^Fcgr4^high^ and Ly6C^high^Fcgr4^low^ in CD11b+NK1.1-Ly6G cell populations were sorted as Ly6C^low^ and Ly6C^high^ monocytes, respectively. Cell sorting was performed using a FACS ARIA III cell sorter (BD Biosciences).

### RNA-seq analysis

Ly6C^low^ and Ly6C^high^ splenic monocytes were obtained by FACS sorting from WT, *Slc29a3*^−/−^, and *Slc29a3*^−/−^ *Tlr7*^−/−^ mice. Total RNA was extracted using RNeasy Mini Kits (Qiagen, Germany), and quality of RNA was evaluated using the Agilent Bioanalyzer device (Agilent Technologies, Santa Clara, CA). The samples with RIN (RNA Integrity Number) value more than 7.3 were subjected to library preparation. RNA-seq libraries were prepared with 1 ng of total RNA using an Ion AmpliSeq Transcriptome Mouse Gene Expression kit (Thermo Fisher Scientific) according to manufacturer’s instructions. The libraries were sequenced on Ion Proton using an Ion PI Hi-Q Sequencing 200 kit and Ion PI Chip v3 (Thermo Fisher Scientific). The FASTQ files were generated using AmpliSeqRNA plug-in v5.2.0.3 in the Torrent Suite Software v5.2.2 (Thermo Fisher Scientific) and analysed by ROSALIND (https://rosalind.bio/, OnRamp Bioinformatics, USA), which is a cloud-based bioinformatics software. Raw reads were trimmed using Cutadapt, and quality scores were assessed using FastQC2. Reads were aligned to the *Mus musculus* genome build mm10 using the STAR aligner. Individual sample reads were quantified using HTseq and normalised via relative log expression (RLE) using the DESeq2 R library. DEseq2 was used to determine the fold changes and *p*-values. Genes showing more than a 1.5-fold change in expression (*p* < 0.05) were considered to be significantly altered. To interpret gene expression profiles, gene set enrichment analysis (GSEA) was performed using MSigDB hallmark gene sets to explore the pathways associated with SLC29A3 deficiency. Enriched pathways with FDR-adjusted *p*-values lower than 0.05 are shown in Figure 2e.

### EdU proliferation assay

*In vivo* and *in vitro* proliferation assays were performed using a Click-iT Plus EdU Alexa Fluor 488 Flow Cytometry Assay Kit (Invitrogen) according to the manufacturer’s instructions. In brief, mice were injected intravenously with 1 mg 5-ethynyl-2’-deoxyuridine (EdU) dissolved in 1xPBS. Spleen and blood samples were collected at 3 h or 3 days after injection. Then, erythrocytes were completely lysed by BD Pharm Lyse lysing buffer (BD Biosciences) to collect splenocytes and peripheral blood mononuclear cells (PBMCs). After blocking splenocytes and PBMCs with anti-CD16/32 (clone:95) mAb, the samples were stained with fluorescent dye-conjugated mAbs. The stained samples were then fixed with BD Cytofix (BD Biosciences) and permeabilised using 1× Click-iT saponin-based permeabilisation and washing reagent. Finally, EdU incorporated into the genomic DNA was stained using Click-iT EdU reaction cocktails. EdU-positive cells were detected using a BD LSR Fortessa cell analyser (BD Biosciences, USA) or a spectral flow cytometer ID7000 (Sony Biotechnology).

### Proliferation assay *in vitro*

Proliferation assays were performed in serum-free AIM-V medium (Thermo Fisher Scientific) supplemented with penicillin-streptomycin-glutamine (Thermo Fisher Scientific). Whole mouse splenocytes and human PBMCs were plated at a density of 5×10^6^ cells per well in a Cepallet W-type 24 well microplate (DIC, Japan) and cultured for 4 days with or without mouse/human M-CSF (Peprotech, USA). Surviving macrophages adhered to 24-well plates were detached by lowering the temperature on ice. The collected cells were incubated with both LIVE/DEAD fixable aqua fluorescent reactive dye (Invitrogen, USA) and SYTOX Green dead cell stain (Invitrogen, USA) in 1×PBS for dead cell staining. Whole mouse splenocytes and human PBMCs were stained with CD11b/Ly6G/NK1.1/Ly6C/Fcgr4 and CD11b/HLA-DR/CD14/CD16 after blocking antibody treatment, respectively. The number of live macrophages was estimated using flow-count fluorospheres (Beckman Coulter, USA) and flow cytometry.

Sorted Ly6C^low^ and Ly6C^high^ monocytes were plated on 96-well plates (BD Falcon, USA) at 2 × 10^4^ cells/well and cultured for 4 days with or without mouse M-CSF. The surviving macrophages were detected by the CellTiter-Glo 2.0 Cell Viability Assay (Promega, USA) following the manufacturer’s protocol, and macrophage number was estimated by comparing with FACS analyses to the control samples whose monocyte numbers were counted.

### LC-MS analysis

Quantitative nucleoside analysis was performed using an LC-MS system equipped with a reversed-phase column (2.0 mm I.D. × 100 mmL) packed with Develosil C30 UG (3μm particle, Nomura Chemical) connected to a hybrid quadrupole-orbitrap mass spectrometer (Q Exactive, Thermo Fisher Scientific) through an electrospray interface. For analyses of nucleoside accumulation in mouse cells and human PBMC-derived macrophages, 1 × 10^7^ cells were lysed using 400 μL D solution (7M guanidine hydrochloride and 0.5M Tris-HCl/10mM EDTANa2, pH 8.5) containing stable isotope-labelled nucleosides (final standard nucleoside concentration; 1 nmol/400 μL of A/U/G/C/dG). For the analysis of nucleoside accumulation in tissues, 100 mg of each tissue was lysed with 400 μL D solution containing stable isotope-labelled nucleosides (final standard nucleoside concentration: 1 nmol/400 μL of C/dG, 10 nmol/400 μL of U/G, and 100 nmol/400 μL of A). The extract was centrifuged at 10,000 × g for 30 min, and the supernatant was diluted 40- to 200-fold with 10 mM ammonium acetate buffer (pH 6.0). Samples (ca. 1–100 pmol nucleosides/40 μL) were loaded into a reversed-phase column and eluted with a 30 min linear gradient from 2% to 12% acetonitrile in 10 mM ammonium acetate buffer (pH 6.0) at a flow rate of 100 μL/min. The eluate from the first 6 min was automatically wasted by switching a 3-way electric valve to remove guanidine hydrochloride from the system and was subsequently sprayed into a mass spectrometer at 3.0 kV operating in positive-ion mode. Mass spectra were acquired at a resolution of 35,000 from m/z 200 to 305. Each nucleoside in the sample cells or tissues was quantified from the peak height relative to that of the corresponding isotope-labelled standard nucleoside. All LC-MS data were processed and analysed using Xcalibur (version 3.0.63, Thermo) and Excel 2013 (Microsoft).

### Platelet and cell counts

Platelet numbers in PBMCs were analysed using an automatic haematology analyser (Celltac α; Nihon Kohden). Cell number was measured using an automated cell counter, CellDrop BF (DeNovix).

### Preparation of splenic B cells

Splenic B cells were purified by negative selection using CD43 MicroBeads (Miltenyi Biotec). Splenocytes from WT and Slc29A3^−/−^ mice were labelled with CD43 magnetic beads, and CD43-negative splenic B cells were enriched using autoMACS (Miltenyi Biotec, Germany) and subjected to experiments.

### Preparation of bone marrow (BM)-derived macrophages and pDCs

BM cells were collected from the tibiae, femora, and pelves of WT, *Slc29a3*^−/−^, and *Slc29a3*^−/−^*Tlr7*^−/−^ mice, and red blood cells were removed using BD Pharm Lyse lysing buffer. For preparation of BM-macrophages (BM-Mphs), BM cells were plated at a density of 7 × 10^6^ cells per well on a non-tissue culture polystyrene 94 mm petri dish (Greiner Bio-One, Germany) and cultured in 10 mL RPMI medium (Gibco™ ThermoFisher Scientific, USA) supplemented with 10% FBS, penicillin-streptomycin-glutamine (Gibco™ ThermoFisher Scientific, USA), 50 μM 2-ME, and 100 ng/mL recombinant murine macrophage colony stimulating factor (M-CSF; PeproTech Inc., USA) for 6 days. Attached cells on petri dishes were collected and used as BM-Mphs. For BM-plasmacytoid DCs (BM-pDCs), BM cells were plated at a density of 2.5 × 10^7^ cells per 10-cm cell culture dish (Greiner Bio-One, Germany) and cultured in 10 mL RPMI 1640 medium (Gibco, Paisley, UK) supplemented with 10% FBS, penicillin-streptomycin-glutamine, 50 μM 2-ME, and 100 ng/mL recombinant murine FMS-like tyrosine kinase-3 ligand (Flt3L, PeproTech Inc., USA) for 7 days. Flt3L-induced pDCs were stained with anti-CD11c/B220 mAbs, and CD11c^+^B220^+^ cells were sorted as BM-pDCs using a FACSAria flow cytometer (BD Biosciences, USA).

### Preparation of human PBMCs and macrophages

All experiments using human samples were approved by the Institutional Ethics Review Boards of the Institute of Medical Science at the University of Tokyo (IMSUT), Jichi Medical University, and Hiroshima University.

To prepare human peripheral blood mononuclear cells (hPBMCs), 7 mL of EDTA-anticoagulated whole blood was treated with 45 mL BD Pharm lysis buffer (BD Biosciences) to completely lyse red blood cells. hPBMCs collected after centrifugation were subjected to FACS analysis and survival assays or allowed to differentiate into macrophages. To induce human macrophages, hPBMCs were plated in 94 × 16 mm petri dishes (Greiner, Germany) at a density of 1.0 × 10^7^ cells per dish and cultured in 10 mL RPMI 1640 medium (Gibco, UK) supplemented with 10% FBS, penicillin-streptomycin-glutamine (Gibco, UK), 50 μM 2-ME, 100 ng/mL of recombinant human M-CSF (PeproTech Inc., USA), and 20 ng of recombinant human IL-4 (PeproTech Inc., USA) for 7 days. After removing floating cells with 1xPBS, the attached cells were collected as human macrophages and subjected to LC-MS and ELISA.

### Cytokine measurements by ELISA

Mouse thioglycollate-elicited PECs and mouse BM-Mphs were cultured in flat-bottom 96-well plates (BD Falcon, USA) at 1 × 10^5^ cells/well. Human PBMC-derived macrophages were cultured in flat-bottomed 96-well plates at 1 × 10^4^ cells/well. All types of immune cells were stimulated with the indicated ligands for 16–20 h, and cytokine concentrations in the supernatant were measured using ELISA. Serum cytokine concentrations in the mice were measured using ELISA. The concentrations of mouse IL-12p40, mouse TNF-α, mouse IL-1β, mouse IL-6, and human TNF-α in the supernatant were measured using Ready-Set-Go! ELISA kits (eBioscience, USA). Mouse IFN-α and IFN-β concentrations in the supernatant were measured using IFN-α/β ELISA kits (PBL Assay Science, USA).

### Cytokine antibody array

The production of 111 cytokines by splenic monocytes was quantified using the Proteome Profiler Mouse XL Cytokine Array (R&D Systems). Cytokine antibody array was performed according to the manufacturer’s instructions. In brief, the antibody array membrane was incubated with 200 μL culture supernatant of splenic monocytes overnight at 4 °C. After incubation with the samples, the membranes were sequentially treated with a detection antibody cocktail and streptavidin-HRP. Finally, the membranes were treated with ECL Select Western Blotting Detection Reagent (GE Healthcare), and the chemiluminescent signal on the membranes was detected using an ImageQuant LAS 500 imager system (GE Healthcare). The intensity of each spot was quantified using the Quick Spots image analysis software (Western Vision Software).

After the cytokine antibody array, IL-6 production by the splenic monocyte population was further determined by flow cytometry to confirm the results from the Proteome Profiler Antibody Arrays.

### Cell death induction and phagocytosis assay *in vivo*

Cell death was induced by treatment of thymocytes at 47 °C for 20 min, and then, cells were incubated at 37 °C for 3 h before subjecting the cell corpses to the phagocytosis assay. Before the phagocytosis assay *in vivo*, cell corpses were stained with the PKH26 Red Fluorescent Cell Linker Kit for General Cell Membrane Labelling (Sigma-Aldrich) according to the manufacturer’s instructions. Then, 5 × 10^7^ PKH26 stained cell corpses were intravenously administered to mice, and mouse spleens were collected 2 h after cell corpse administration. Immune cells engulfing PKH26-positive cell corpses were detected using flow cytometry.

### Lentiviral transduction

FLAG-tagged human SLC29A3 was expressed in the mouse macrophage cell line J774.1 cells using lentiviral transduction. The cDNA of FLAG-SLC29A3 was substituted with the BFP2Apuro sequence in the lentiviral pKLV-U6gRNA(BbsI)-PGKpuro2ABFP vector (Addgene plasmid 50946), excluding the U6gRNA(BbsI) site. The ViraPower Lentiviral expression system (Thermo Fisher Scientific) was used to prepare the lentivirus for Flag-SLC29A3 overexpression according to the manufacturer’s instructions. Supernatants containing lentivirus particles were collected 24 h post-transfection and used for transduction.

### Structured illumination microscopy

Macrophages from the J774.1 cell line were allowed to adhere to collagen-coated coverslips overnight and were stimulated by 1 mM Phorbol 12-myristate 13-acetate (PMA) for 2 h. Cells attached on coverslips were treated with the heat treated dying thymocytes for 1 h. After engulfment, the cells were fixed with 4% paraformaldehyde for 10 min and then permeabilized with 1XPBS containing 0.2% saponin for 30 min.

After blocking with 2.5% BSA Blocking One (Nacalai Tesque, Japan) for 30 min, cells were incubated with anti-TLR7 antibody and anti-HA antibody (Roche) at 37 °C for 90 min. After washing the cells three times, the cells were incubated for 90 min at 37 °C with AlexaFluor-488 conjugated goat anti-mouse and AlexaFluor-568 goat anti-rat antibodies and DAPI (Invitrogen). Fluorescence microscopy was performed using a Nikon Structured illumination microscope (N-SIM, Nikon) at excitation wavelengths of 405, 488, and 561 nm with a CFI Apochromat TIRF 100 × objective lens (1.49 NA, NIKON). Data acquisition was performed in 3D SIM mode before image reconstruction using NIS-Element software. Each image represents more than three independent experiments.

### Statistical analysis

Statistical significance between the two groups was determined using a two-tailed, unpaired t-test with Holm–Sidak correction. To determine significant differences between more than three groups, one-way ANOVA followed by Dunnett’s multiple comparison test was employed in this study. All data are represented as the mean ± standard deviation (s.d.) and graphs were made using PRISM. Statistical significance was set at p < 0.05. \**p* < 0.05, \*\**p* < 0.01, \*\*\**p* < 0.001.

## Acknowledgements

We thank Prof. P. W. Kincade for critically reviewing the manuscript and Dr. Xiaobing Li for supporting our work. We thank Dr. Shota Endo for kindly providing *Hcst*^−/−^ mouse embryos and Mrs. Noriko Tokai for helping us analyse the imaging results. We acknowledge Dr. Haruya Ohno and Dr. Masayasu Yoneda at Hiroshima University for providing the patient sample. We would like to thank Editage (www.editage.com) for English language editing. This work was supported in part by JSPS/MEXT KAKENHI Grants: 16H06388, 21H04800, 22H05184, 22K19424, and JP22H05182 to K. M; 16H02494 to T. Shimizu; 21K15464 to R. S.; 26293083 to S.-I.S.; 19H03451 and 16K08827 to T.S.; and JST CREST (JPMJCR13M5, JPMJCR21E4) to T. Shimizu.; the Japan Agency for Medical Research and Development (AMED) Grant Number JP20ek0109385 to T.S.; The Mochida Memorial Foundation for Medical and Pharmaceutical Research to T.S.; Joint Research Project of the Institute of Medical Science at the University of Tokyo; and JSPS KAKENHI Grant Number JP 16H06276 (AdAMS).

## Contributions

T.S. and K.M. conceived of and designed the experiments. T.S., M.O., and N.Y. constructed knockout and transgenic mice. M.T., Y.Y. and T. Isobe conducted LC-MS analysis. R.S. and S.S. performed high-resolution microscopy. T.S., E.F., M.K, M.O., E.L., and A.T. analysed human blood samples from patients with H syndrome. K.Y. and F.Y. performed the transcriptome analyses. T.S. performed the *in vivo* analyses with the help of S. G., Y. M., J. K., K. I., Y. T., T. Ichinohe, R. H., R. F., T. K., and T. T. R.S., K. Mukai, and T.T. performed imaging and biochemical analyses. T.S. performed all other experiments in this study and analysed the data for all figures. T.S. and K.M. wrote the paper with assistance from U.O., and T. Shimizu. All authors have reviewed the manuscript.

## Ethics declarations

competing interests

This work was partially funded by Daiichi Sankyo Co., Ltd.

## Supplementary Materials

**Extended Data Figure 1.**
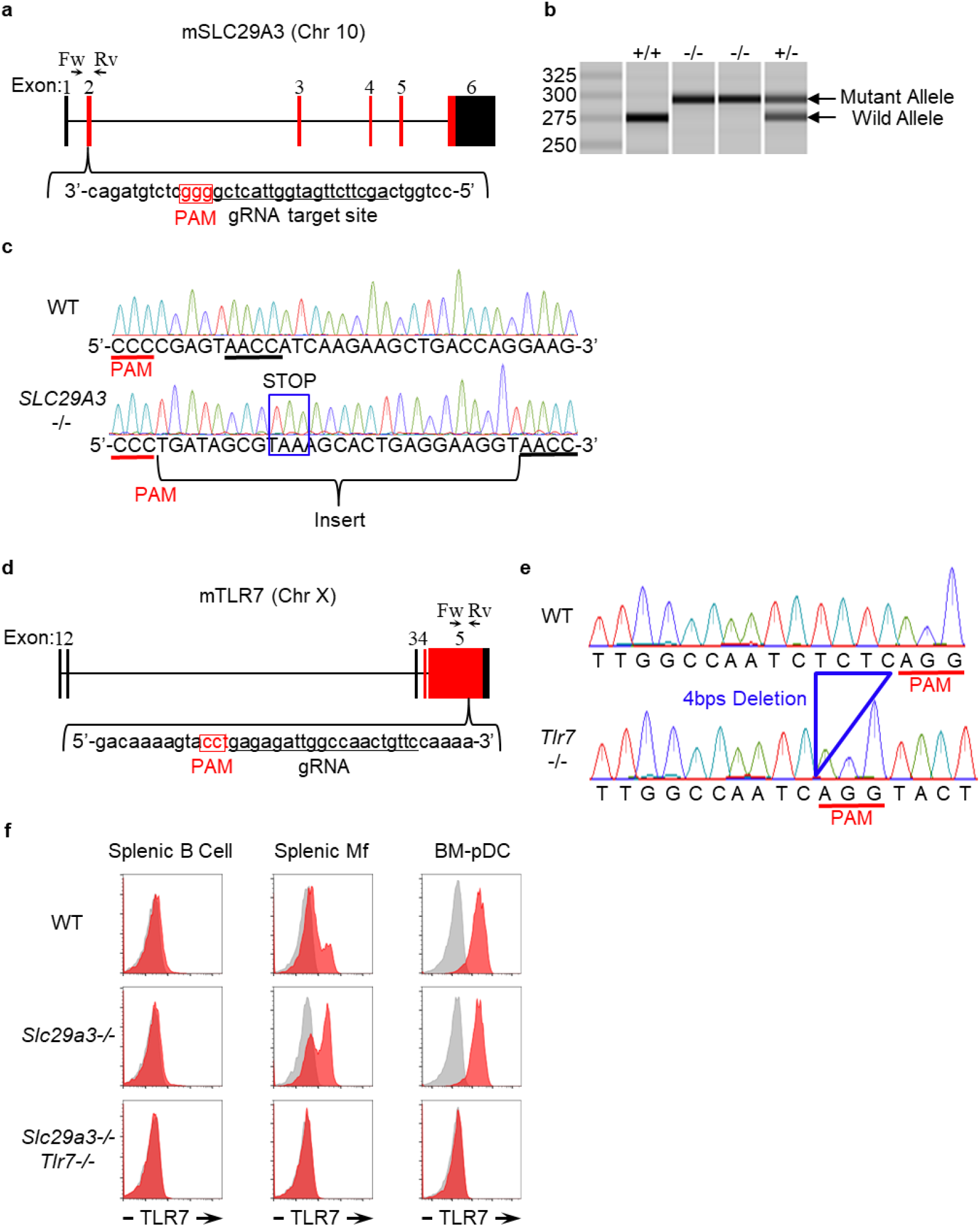
Generation of *Slc29a3*^−/−^ and *Tlr7*^−/−^ mice. **a, d,** Genomic configuration of *Slc29a3* and *Tlr7* genes showing 20mer gRNA target sites to introduce a mutation into exon 2 of *Slc29a3* and exon 5 of *Tlr7*. The PAM sequence is highlighted by the red box. **b**, Genomic PCR with the primer set (Fw and Rv) shown in (**a**) to reveal an insertional mutation in the targeted allele of *Slc29a3*. **c**, Direct sequencing of the gRNA target site of *Slc29a3*. The inserted sequence containing the stop codon is shown by the blue box. **e**, Sequence data of the gRNA target site on the *Tlr7* allele showing a 4-bp deletion in the 5^th^ exon of *Tlr7* (blue). **f**, FACS analysis showing the lack of TLR7 protein in splenic B cells, splenic monocytes, and BM-derived plasmacytoid DCs from WT, *Slc29a3*^−/−^, and *Slc29a3*^−/−^ *Tlr7*^−/−^ mice. Red and grey histograms represent intracellular staining with and without anti-TLR7 mAb, respectively.

**Extended Data Figure 2.**
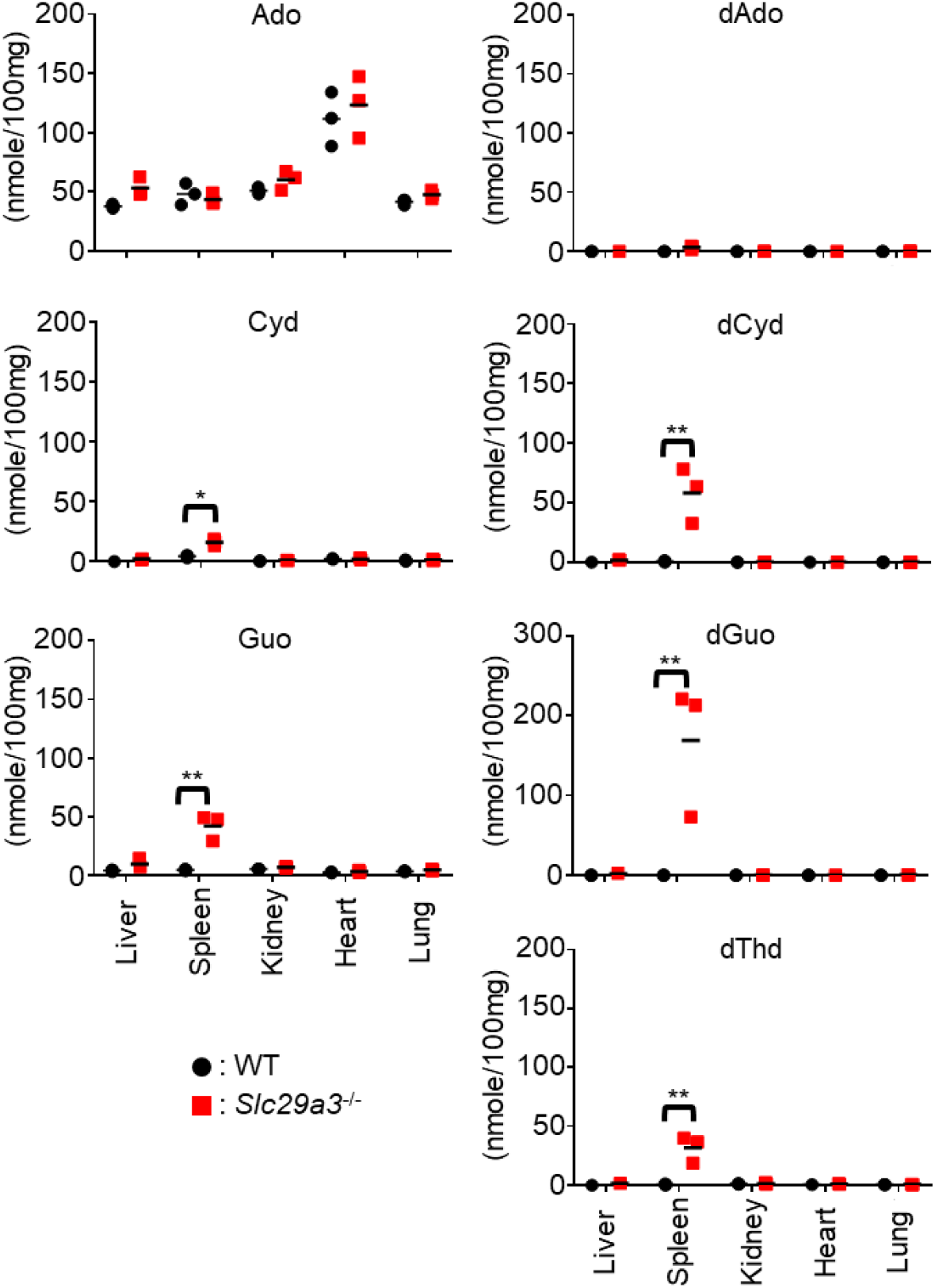
Nucleoside storage in *Slc29a3*^−/−^ mice. The amount of nucleosides (nmol/100 mg) in each organ from WT and *Slc29a3*^−/−^mice was determined by LC-MS analyses. Dot plots represent values from the indicated mice (n = 3). \**p* < 0.05, \*\**p* < 0.01.

**Extended Data Figure 3.**
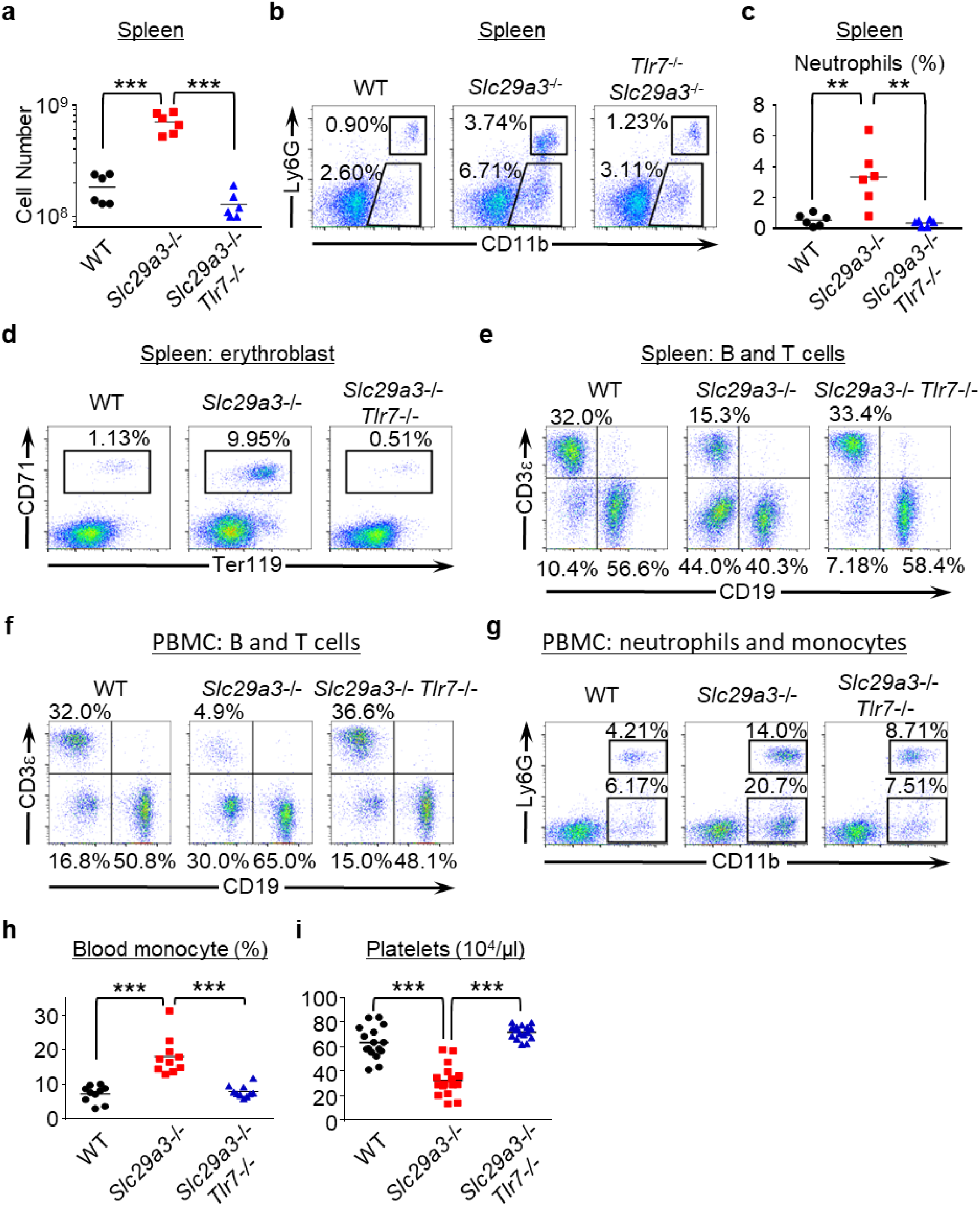
TLR7-dependent monocytosis in *Slc29a3*^−/−^ mice. **a**, Dot plots showing the number of splenocytes in the indicated mice (n = 6). **b**, Representative FACS analyses of Siglec F^-^ NK1.1^-^ splenocytes to show their expression of CD11b and Ly6G. **c**, Each dot shows the percentage of neutrophils in whole splenocytes from the indicated mice (n = 6). **d**, **e**, FACS analyses of CD71^+^ Ter119^+^erythroblasts (**d**) or CD19^+^B and CD3ε^+^ T cells (**e**) in the spleen. **f**, **g**, FACS analyses of B and T cells (**f**) or neutrophils and monocytes (**g**) in PBMCs. **h**, **i**, Percentages of CD11b^+^ Ly6G^-^ monocytes (n= 10) in PBMCs (**h**) and platelet counts (n = 16) in the peripheral blood (**i**) of 3-month-old mice.

**Extended Data Figure 4.**
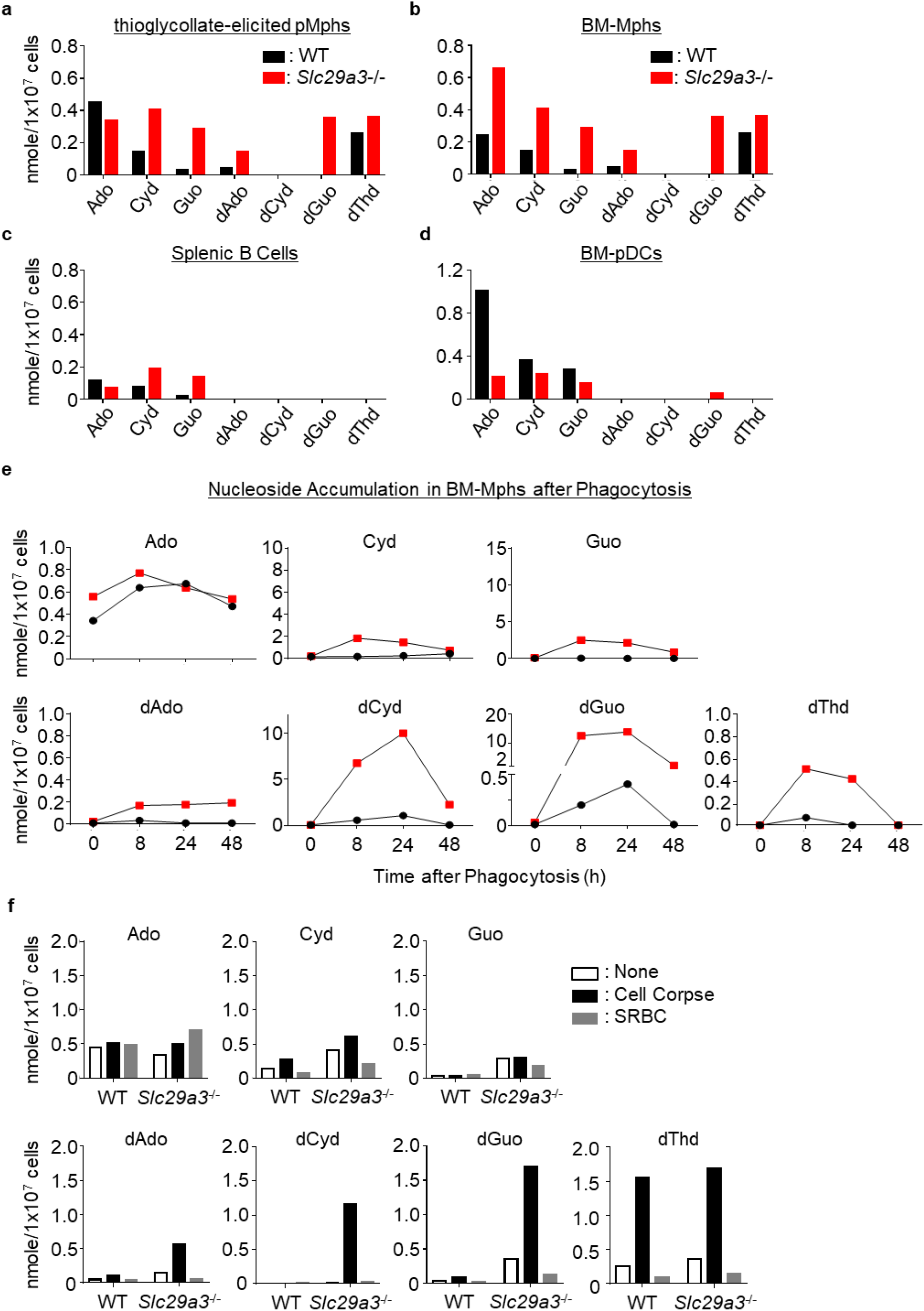
Nucleoside storage in *Slc29a3*^−/−^ phagocytes. **a-d**, The amount of nucleosides (nmol) in 10^7^ thioglycolate-elicited pMphs (**a**), bone marrow-derived macrophages (BM-Mphs) (**b**), splenic B cells (**c**), and BM-plasmacytoid dendritic cells (BM-pDCs) (**d**) from WT (black) and *Slc29a3*^−/−^ (red) mice. **e**, The amount of nucleosides (nanomoles) in 10^7^ BM-Mphs of WT and *Slc29a3*^−/−^ mice at the indicated time points after treatment with 10^8^ dying thymocytes (cell corpse). **f**, The amount of nucleosides (nanomoles) in 10^7^ BM-Mphs of WT and *Slc29a3*^−/−^ mice after 1 day treatment with 10^8^ dying thymocytes or 10^9^ SRBC.

**Extended Data Figure 5.**
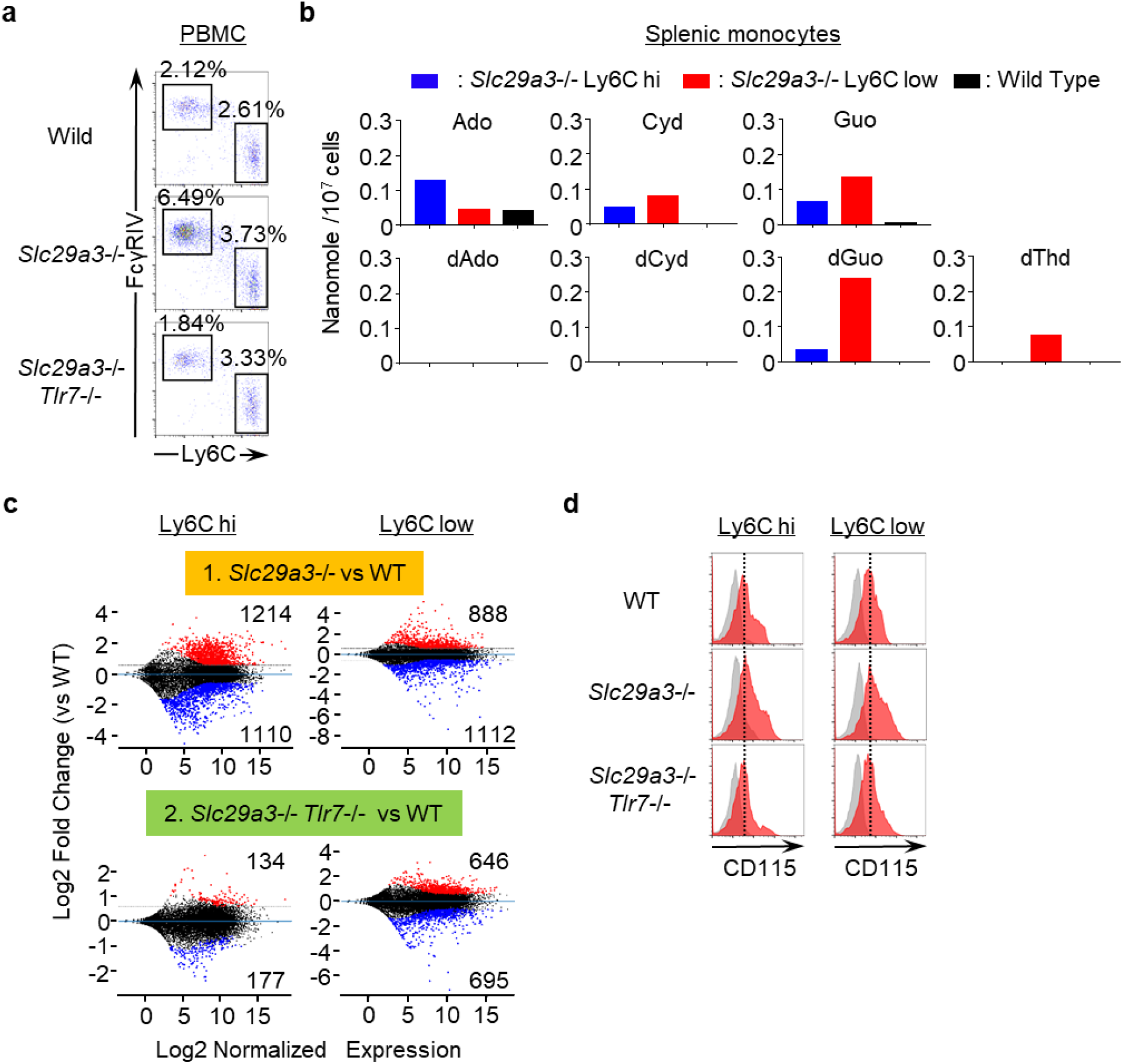
Analyses of Ly6C^hi^ and Ly6C^low^ monocytes. **a**, Representative FACS analyses showing the expression of FcγRIV and Ly6C on CD11b^+^ Ly6G^-^ monocytes in the peripheral blood. The percentages of FcγRIV^hi^ Ly6C^low^ patrolling and FcγRIV^low^ Ly6C^hi^ classical monocytes among whole peripheral mononuclear cells are shown. **b**, The amount of nucleosides (nmole/10^7^ cells) in *Slc29a3*^−/−^ Ly6C^hi^ and Ly6C^low^ splenic monocytes or in WT CD11b^+^ splenic monocytes. **c**, Transcriptome analyses of Ly6C^hi^ and Ly6C^low^ monocytes from the spleen. MA-plots displaying Log2 normalized expression (X-axes) and Log2 fold change of expression (Y-axes) for the comparisons of *Slc29a3*^−/−^ (n = 4) vs WT (n = 4) monocytes and *Slc29a3*^−/−^ *Tlr7*^−/−^ (n = 4) vs WT (n = 4) monocytes. More than 1.5-fold upregulated and downregulated genes are shown in red and blue, respectively. **d**, Red histograms show cell surface expression of CD115 in Ly6C^hi^ and Ly6C^low^ monocytes in the spleens of WT, *Slc29a3*^−/−^, and *Slc29a3*^−/−^ *Tlr7*^−/−^ mice. Grey histograms show staining with control mAb.

**Extended Data Figure 6.**
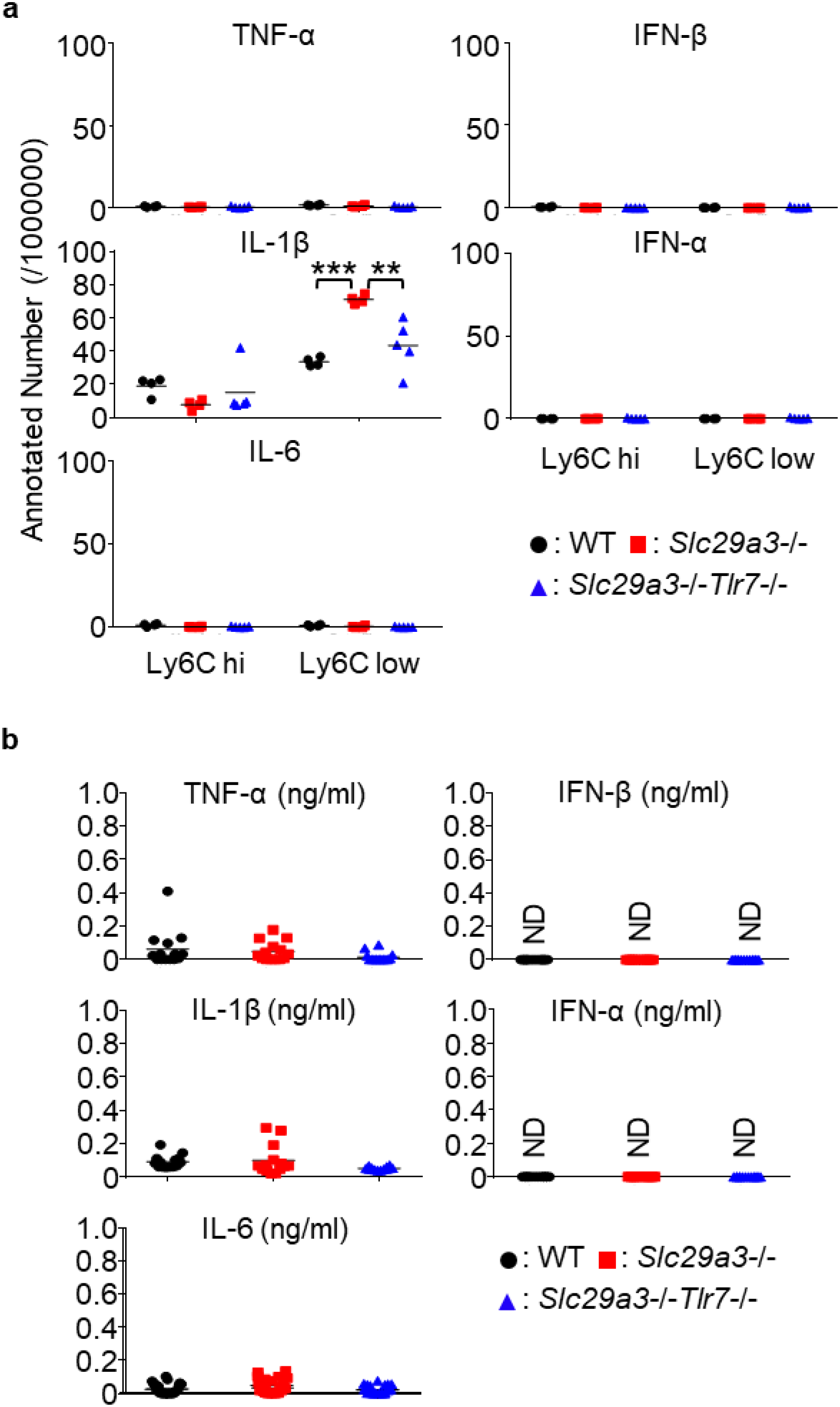
Low cytokine production in *Slc29a3*^−/−^ mice. **a**, mRNA expression of cytokines in splenic Ly6C^hi^ and Ly6C^low^ monocytes from WT (black), *Slc29a3*^−/−^ (red), and *Slc29a3*^−/−^ *TLR7*^−/−^ (blue) mice. Each dot shows the normalised read count per 1 million reads from the RNA-seq analyses for each monocyte population (n = 4). **b**, Serum cytokine levels were determined using ELISA. Sera were collected from 4–6 months old mice (n = 14). \*\**p* < 0.01, \*\*\**p* < 0.001. ND, not detected.

**Extended Data Figure 7.**
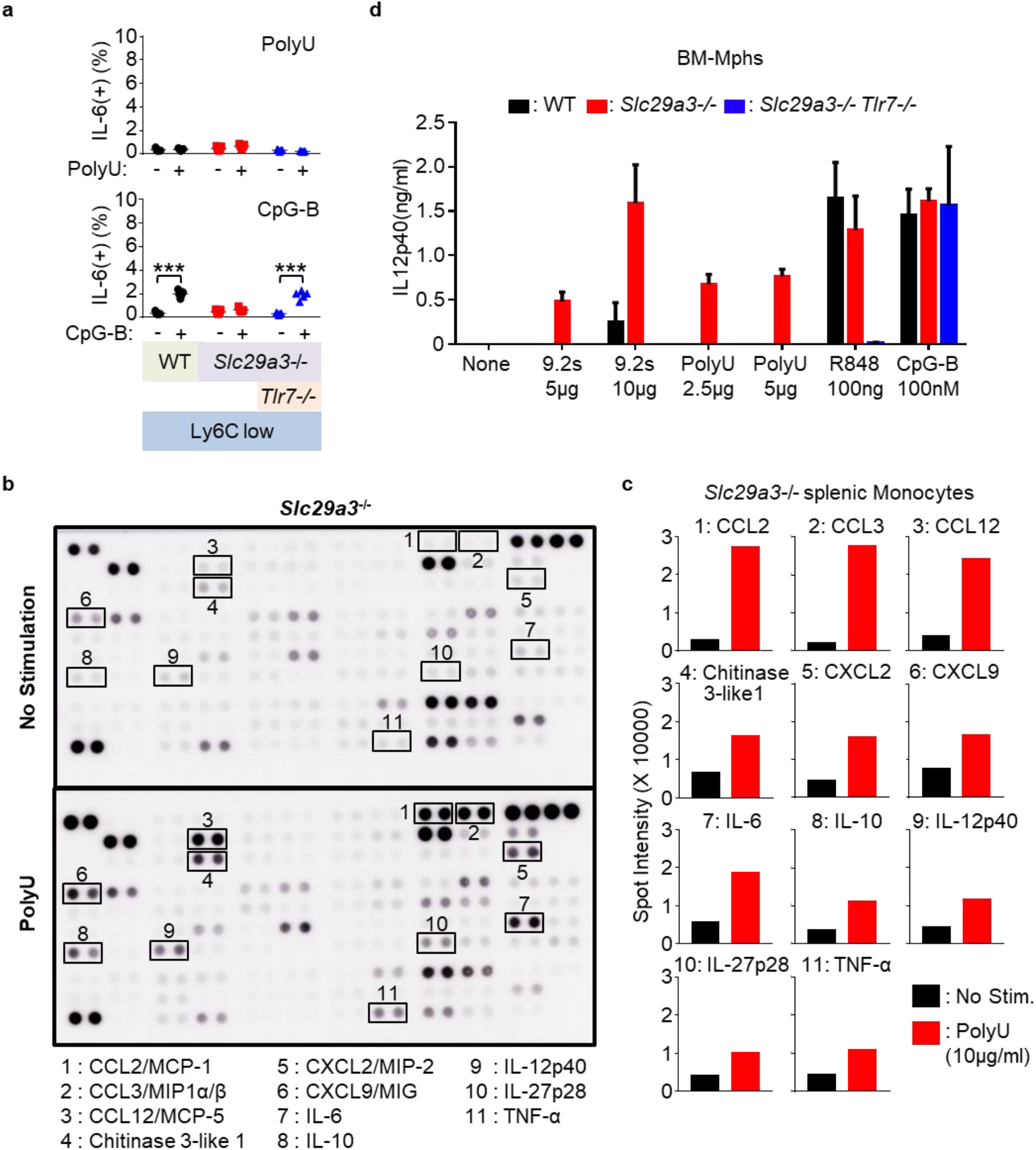
Enhanced TLR7 response to ssRNA in *Slc29a3*^−/−^ monocytes. **a**, The percentage of IL-6^+^ cells in Ly6C^low^ monocytes from indicated mice after *in vitro* stimulation with poly U (10 μg/mL) or CpG-B (1 μM) in the presence of brefeldin A (10 μg/mL) for 4 h. Each dot represents the value for each mouse (n = 5). **b, c**, *Slc29a3*^−/−^ splenic monocytes were stimulated with poly U (10 μg/mL) for 18 h. Cytokines in the supernatants were detected using a cytokine antibody array (**b**). The results are shown as the mean signal intensity value for each cytokine spot (**c**). **d**, IL-12 p40 production by BM-Mphs after stimulation with TLR7 and TLR9 ligands for 18 h. The results are represented as the mean ±s.d. from triplicate samples.

**Extended Data Figure 8.**
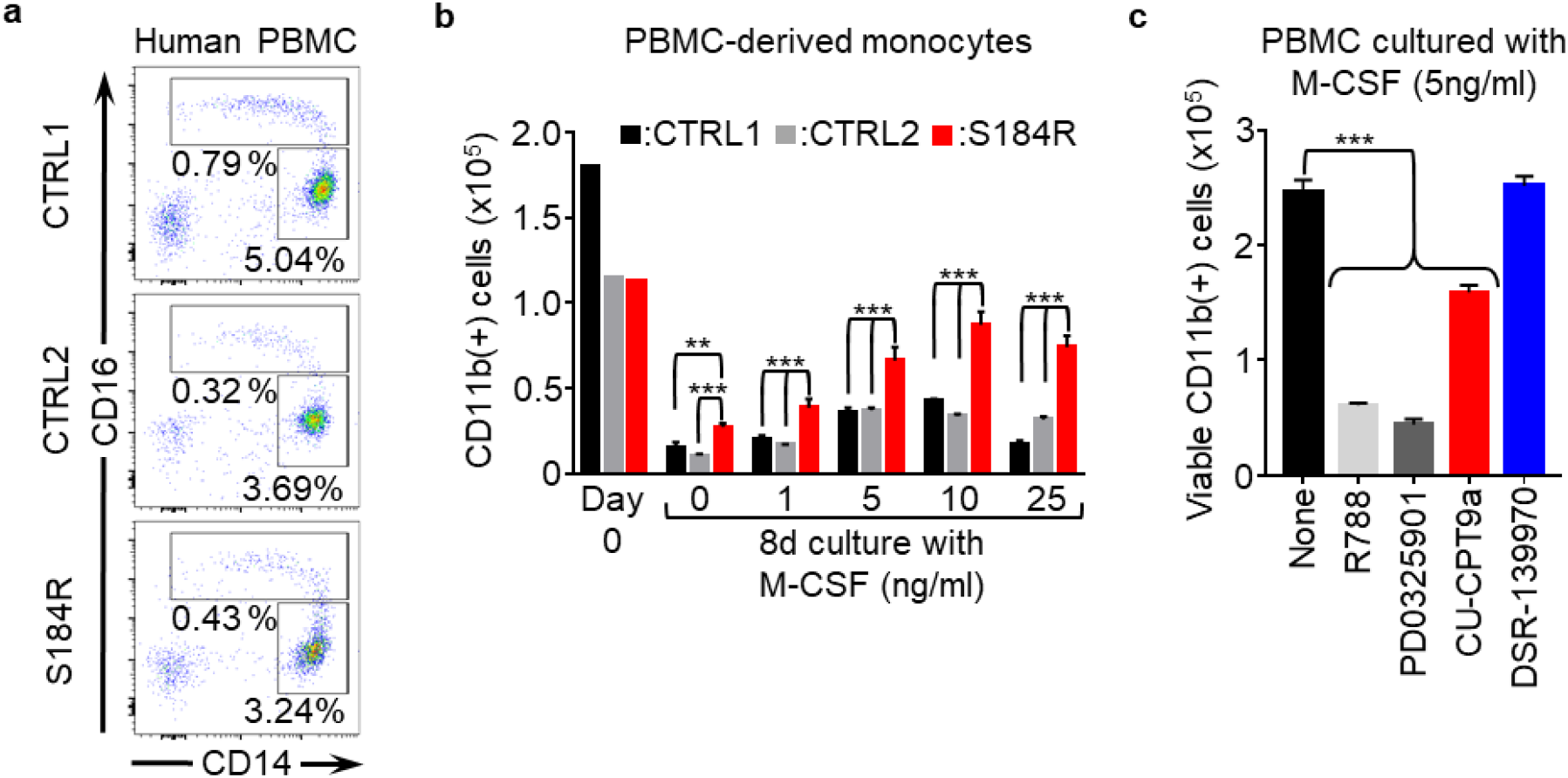
Enhanced survival of PBMC derived monocytes from the patient harbouring the S184R *SLC29A3* mutation. **a**, Expression of surface CD16/CD14 in HLA-DR^+^ CD15^-^ CD56^-^ PBMCs from the patient with the S184R *SLC29A3* mutation and from healthy subjects. **b**, Numbers of CD11b^+^ CD15^-^ CD56^-^ monocytes in PBMCs from the patients (red) and healthy subjects (black and grey) that survived 4 days of culture with M-CSF at the indicated concentrations. **c,** Numbers of CD11b^+^ cells that survived 8 days PBMC culture with 5 ng/mL M-CSF in the absence or presence of the indicated inhibitors of Syk (1 μM R788), MEK1/2 (1 μM PD0325901), TLR8 (10 μM CPT9a), and TLR7 (10 μM DSR-139970).

